# Growth-dependent concentration gradient of the oscillating Min system in *Escherichia coli*

**DOI:** 10.1101/2023.08.01.551406

**Authors:** Claudia Parada, Ching-Cher Sanders Yan, Cheng-Yu Hung, I-Ping Tu, Chao-Ping Hsu, Yu-Ling Shih

## Abstract

The Min system contributes to the spatiotemporal regulation of division sites in *Escherichia coli*. The MinD and MinE proteins of this system self-organize into oscillatory waves in the form of concentration gradients. How the intracellular Min protein concentration gradients are coordinated with cell growth to achieve spatiotemporal accuracy of cell division is unknown. Here, we report that the MinD concentration gradient becomes progressively steeper as cells elongate, suggesting that the division inhibitory activity at the midcell also decreases with cell growth. Interestingly, the oscillation period appears relatively stable across different cell lengths. Similar features were found in cells under carbon stress conditions, but the gradient was even steeper, likely favoring division at shorter cell lengths. The length-dependent variation of the concentration gradient was further examined *in silico* using a reaction-diffusion model, which not only supported the above features, but also revealed a decrease in the midcell concentration as the shape of the gradient becomes steeper in growing cells. This growth-dependent regulation of the midcell concentration of MinD may be coupled with the FtsZ ring formation through the MinD-interacting protein MinC. We found that the variable concentration gradients occur by coordinating the reaction rates of the recruitment of MinD and MinE to the membrane and the recharging of MinD with ATP in the cytoplasm. In conclusion, this work uncovers the plasticity of MinD concentration gradients during interpolar oscillations throughout cell growth, an intrinsic property integrated during cell division.

## Introduction

Self-organization provides spatiotemporal information about the cellular organization of biological processes. This self-organizing phenomenon of proteins is exemplified by the the Min system of *Escherichia coli*, which undergo spontaneous oscillations that mediate the placement of the division septum at the midcell. Although a large number of biochemical and biophysical studies are available to understand the complex molecular interactions behind the formation of the oscillatory modes of MinD and MinE (Denk *et al*, 2018; Lutkenhaus, 2007; Shih & Zheng, 2013; Vecchiarelli *et al*, 2016), mapping *in vitro* features and molecular interactions back to the cellular environment remains a formidable challenge. This is partly due to differences in spatial and temporal scales and the reduced complexity of *in vitro* reconstitution experiments. Interest in the *in vivo* versus *in vitro* characterization of the ‘Min system and its relationship to overall bacterial physiology has prompted further investigations of Min protein oscillations in the cellular context.

MinD is a deviant form of the Walker-type ATPase, and MinE drives MinD oscillations by stimulating the ATPase activity of MinD (Hu & Lutkenhaus, 2001; Zhou *et al*, 2005). In a cellular context, the oscillation cycle starts with a fully assembled MinD polar zone covering half the cell, and then switches to disassembly upon MinE stimulation. Dissociated MinD molecules diffuse to the opposite cell pole during polar zone disassembly and reassemble into the polar zone until half of the cell is covered. The new polar zone then disassembles, and the molecules diffuse back to the original pole to reassemble into a polar zone, completing an oscillation cycle (Fig. 1A). During oscillation, the division inhibitor MinC interacts with MinD and oscillates, destabilizing the FtsZ polymers at the poles (Hu *et al*, 1999; Raskin & de Boer, 1999a, b), thereby promoting the formation of the FtsZ ring that assembles the dividing septum (de Boer *et al*, 1988; Fu *et al*, 2001; Raskin & de Boer, 1999a, b). Mechanistically, the oscillation cycles of the Min system arise from a balance of concentration ratios between MinD and MinE and their complex interactions with the membrane (Mizuuchi & Vecchiarelli, 2018; Vecchiarelli *et al*., 2016). In the time-averaged format, the concentration of MinD along the long axis of the cell is higher at the poles but lower at the midcell.

**Fig. 1.**
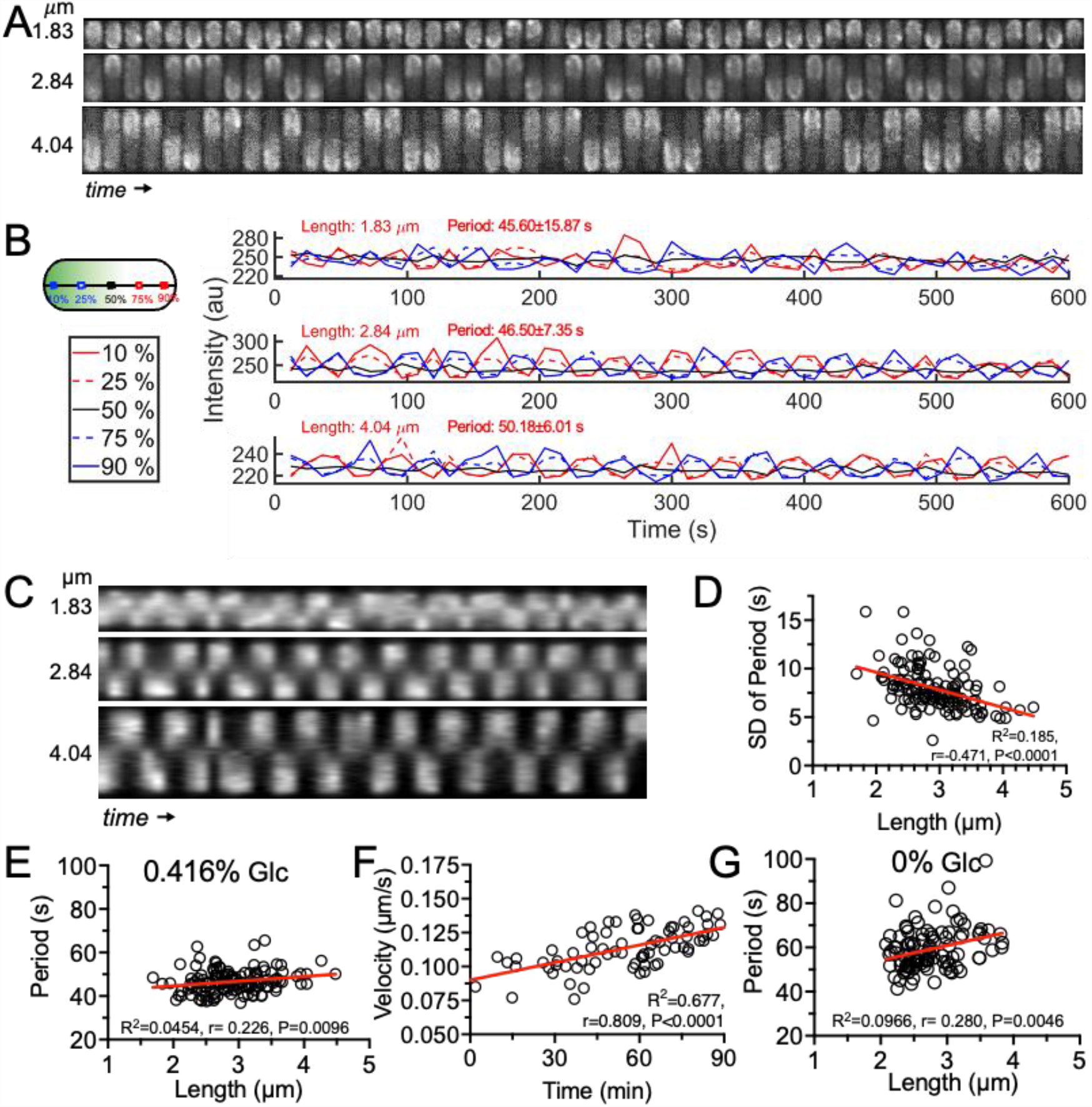
MinD oscillation in cells of different lengths. **A**, Time-lapse micrographs showing interpolar oscillations of sfGFP-MinD in cells of different lengths. Exponentially growing *E. coli* strain FW1541 was imaged for 10 min at 12-sec intervals. **B**, The standing wave features are manifested at 10% and 25% of the cell length, corresponding to 90% and 75% along the cell’s mid-axis. **C**, Demographs showing distribution of intensity as a function of length. **D**, The standard deviation (SD) of the oscillation period decreases in longer cells. **E**,**G**, Correlation plots between oscillation period and cell length when incubated with 0.4% (E) and 0% glucose (G). **F**, Correlation between oscillation velocity and cell cycle time. In D-G, the red line represents the fitted line of simple linear regression. n, population size; R^2^, goodness of fit; r, Spearman coefficient; P: two-tailed probability. D-F, n=130; G, n=101.

A piece of missing information needed to link underlying biochemical and molecular interactions to oscillations in the cellular environment is the spatiotemporal distribution (i.e., the concentration gradient) of Min proteins through the cell cycle. By tracking the shape of the MinD concentration gradient during oscillations, the current study found that the MinD concentration gradient gradually steepens as cells elongate. In addition, the concentration of MinD at the midcell progressively decreases as the cell elongates, and then remains at a lower concentration until the cell divides. The distribution of the division inhibitor MinC may be synchronized with spatiotemporal differences in MinD concentrations, leading to a stable placement of the FtsZ ring at the midcell. Surprisingly, it was found that the oscillation period did not change significantly as cells grow. These *in vivo* observations are supported by a numerical model of a reaction-diffusion system that exhibits a similar coupling between variable concentration gradients and decreasing concentrations in MinD at the midcell. The model also predicts that the MinD concentration gradient can scale proportionally to the length of the cell approaching division. Altogether, these results provide new insights into concentration gradients of the Min system in the cellular environment.

## Results

### Endogenous expression of sfGFP-MinD ensures characteristics of oscillations

This study used exponentially growing *E. coli* cells of the strain FW1541 expressing endogenous levels of sfGFP-minD from the native chromosomal locus (Wu *et al*, 2015) (Table S1). The cellular abundance of sfGFP-MinD and MinE in exponentially growing cells were determined to be 2205±178.3 molecules per cell (1.95±0.16 μM) and 1580±148 molecules per cell (1.4±0.13 μM), respectively. Meanwhile, the cellular abundance of MinD and MinE of the parental strain W3110 were 3532 ± 61.3 molecules per cell (3.12 ± 0.05 μM) and 2150 ± 228 molecules per cell (1.9 ± 0.2 μM) (Fig. S1). Although the expression levels of both sfGFP-MinD and MinE were lower in FW1541, the concentration ratios between MinD and MinE were close (sfGFP-MinD/MinD to MinE: FW1541, 1.40; W3110, 1.64). Therefore, expression of sfGFP-MinD and MinE in FW1541 is comparable to that in the parental strain W3110 and in previous reports (Hale *et al*, 2001; Li *et al*, 2014; Schmidt *et al*, 2016; Shih *et al*, 2002).

Classical features of MinD oscillations, as well as novel ones, were identified by tracking the distribution of fluorescence intensity measured over time along the cell’s medial axis (Fig. 1, S2). First, a standing wave pattern of interpolar oscillations is demonstrated by the registration of sfGFP-MinD intensity profiles located at different positions on opposite cell halves (Fig. 1A, B). While waves from opposite halves of a cell are recorded as being out of phase, waves from different locations on the same half of the cell are recorded as being nearly in phase (Fig. 1B). Second, periodic regularity of interpolar oscillations increases with cell length, as evidenced by kymographs depicting one-dimensional fluorescence intensity over time (Fig. 1C). This is supported by the gradual stabilization of periodicity as cells approach division, and the decrease in standard deviation of the oscillation period with cell length (Fig. 1D). These observations echo previously reported changes in oscillation patterns before and after cell division (Juarez & Margolin, 2010). Third, the oscillation period (defined as the time it takes for MinD to move back and forth between the two cell poles) was determined to be 46 seconds (median, *n*=130; Table 1). The oscillation velocity was calculated to be 0.127 μm/sec by dividing the period by the two cell lengths, the distance that MinD travels in one oscillation cycle. These results are not only comparable to previous studies (Fu *et al*., 2001; Hale *et al*., 2001; Shih *et al*., 2002), but also close to physiological conditions than studies reported overexpression from plasmids (Di Ventura & Sourjik, 2011; Meacci & Kruse, 2005; Tostevin & Howard, 2006; Touhami *et al*, 2006) or under the control of the *lac* promoter on chromosomes (Juarez & Margolin, 2010).

**Table 1.**
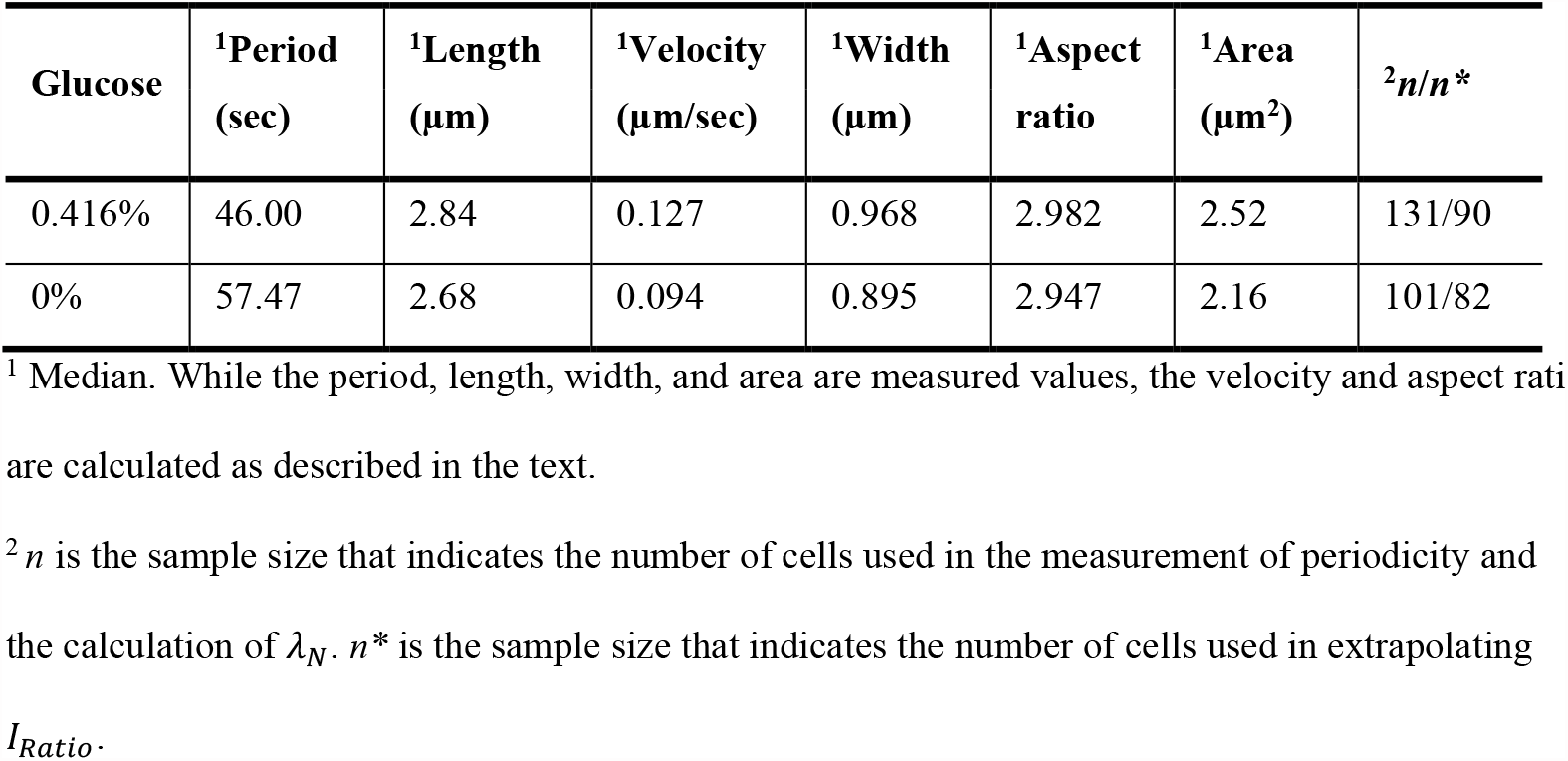
Characteristics of sfGFP-MinD oscillation in strain FW1541 (Wu *et al*., 2015).

| Glucose | <sup>1</sup>Period<br>(sec) | <sup>1</sup>Length<br>($\mu\text{m}$) | <sup>1</sup>Velocity<br>($\mu\text{m}/\text{sec}$) | <sup>1</sup>Width<br>($\mu\text{m}$) | <sup>1</sup>Aspect<br>ratio | <sup>1</sup>Area<br>($\mu\text{m}^2$) | <sup>2</sup>$n/n^*$ |
| --- | --- | --- | --- | --- | --- | --- | --- |
| 0.416% | 46.00 | 2.84 | 0.127 | 0.968 | 2.982 | 2.52 | 131/90 |
| 0% | 57.47 | 2.68 | 0.094 | 0.895 | 2.947 | 2.16 | 101/82 |

<sup>1</sup> Median. While the period, length, width, and area are measured values, the velocity and aspect ratio are calculated as described in the text.
<sup>2</sup> $n$ is the sample size that indicates the number of cells used in the measurement of periodicity and the calculation of $\lambda_N$ . $n^*$ is the sample size that indicates the number of cells used in extrapolating
$I_{\text{Ratio}}$ .

### Oscillation periods remain relatively stable in cells of different lengths

In addition to these features, the oscillation period was found to be relatively stable for different cell lengths (r=0.226; Fig. 1E). If periodicity is to be maintained regardless of increasing length, the velocity of the Min oscillations must increase as the cell cycle progresses. Here, we only consider cell length, since the shape descriptors of length, width and aspect ratio show little or no difference and they are highly correlated (Fig. S3A, B). Furthermore, since cell length is a function of time in an actively growing cell, we can calculate the corresponding time in the cell cycle from the length (Fig. S3C). Therefore, we were able to conclude that the oscillation period remained relatively constant (Fig. S3D) and the velocity increased with time (Fig. 1F) throughout the cell cycle. Furthermore, faster velocities in longer cells and slower velocities in shorter cells suggest that velocities reset to slower velocities after division. By tracking sfGFP-MinD oscillations in individual cells with clear mother-daughter lineages, we confirmed that the velocity of newborn cells was significantly reduced even when growing under adverse growth conditions during imaging (Fig. S4). Nonetheless, the location of the FtsA-labeled division site was independent of cell length and oscillation period (Fig. S5).

### MinD concentrations barely change during the cell cycle

We wondered how cellular MinD and MinE concentrations contribute to maintaining the oscillation period. Bulk fluorescence of sfGFP-MinD in individual cells was measured from each time frame in the time-lapse sequence. As shown in Fig. 2A, while the fluorescence of sfGFP-MinD gradually increased over time, and the intensity change per unit area (μm^2^) was relatively small. All measurements obtained from the complete cell cycle were then identified, aligned and fitted by simple linear regression. Fluorescence intensity was converted to number of molecules by applying 2205 molecules per cell at the midpoint of the experimental doubling time (74.68 min, n=26) on the fitted curve (Fig. 2B). During a 90-minute cell cycle, 1844–2537 molecules are produced per cell, corresponding to concentrations of 0.92–1.26 μM. Interestingly, the pre-division value did not double the post-division value, which could be the result of a balance between *de novo* synthesis and degradation, or a burst of MinD synthesis followed by continued synthesis in growing cells.

**Fig. 2.**
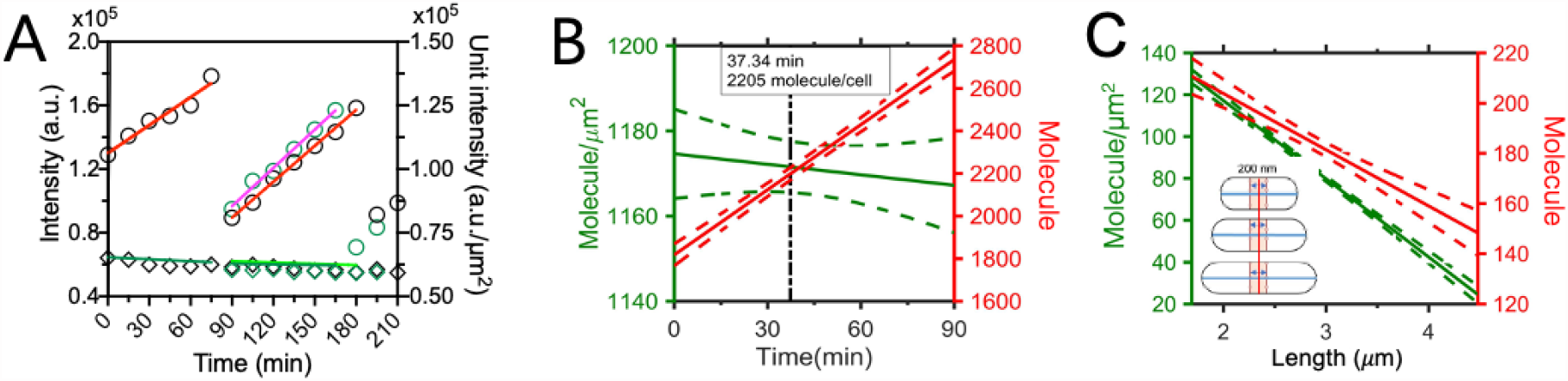
Number of MinD molecules during the cell cycle. **A**. The bulk fluorescence intensity of sfGFP-MinD (circles, red and pink lines) increased over time, but the intensity per unit area (diamonds, green lines) showed only modest fluctuations. The data of paired daughter cells are distinguished by color. The figure contains data from a portion of a time-lapse image sequence acquired at 15-minute intervals over 5 hours. **B**, The number of molecules in a single cell (red line) and the number of molecules per unit area (μm^2^; green line) were plotted against time and fitted by simple linear regression (n = 100). Dashed lines represent 95% confidence intervals. Rapidly growing cells that divide within 5 frames were excluded. **C**, As cells grow longer, the number of molecules within 200 nm of the cell center decreases.

Nonetheless, the number of molecules per square micron fluctuates within a very narrow range. Therefore, the importance of MinD concentration for maintaining the oscillation period may be limited during the cell cycle. However, when focusing on the midcell region defined within 200 nm of the cell midpoint, protein molecules per μm^2^ were reduced to ∼20 upon division (Fig. 2C). This analysis suggests that the spatiotemporal distribution of MinD, i.e. the concentration gradient, may be the key to our problem.

### The wave slope toward the center of the cell becomes steeper as the cell gets longer

The MinD concentration gradient was analyzed in detail (Fig. S2) to understand how it contributes to the aforementioned features of the Min system. Fluorescence intensity profiles of individual cells collected at different time points in the time-lapse image sequence were separated by their position on the left or right pole and fitted by an exponential decay equation. Then, the fitted curves obtained for the entire cell population were compressed into one dimension, sorted by cell length, and compiled into a demographic map to gain a comprehensive understanding of the changes in MinD concentration gradients in cells at different growth stages. The demograph intuitively shows that the MinD concentration (proportional to intensity) near the cell center is higher in shorter cells than in longer cells (Fig. 3A).

**Fig. 3.**
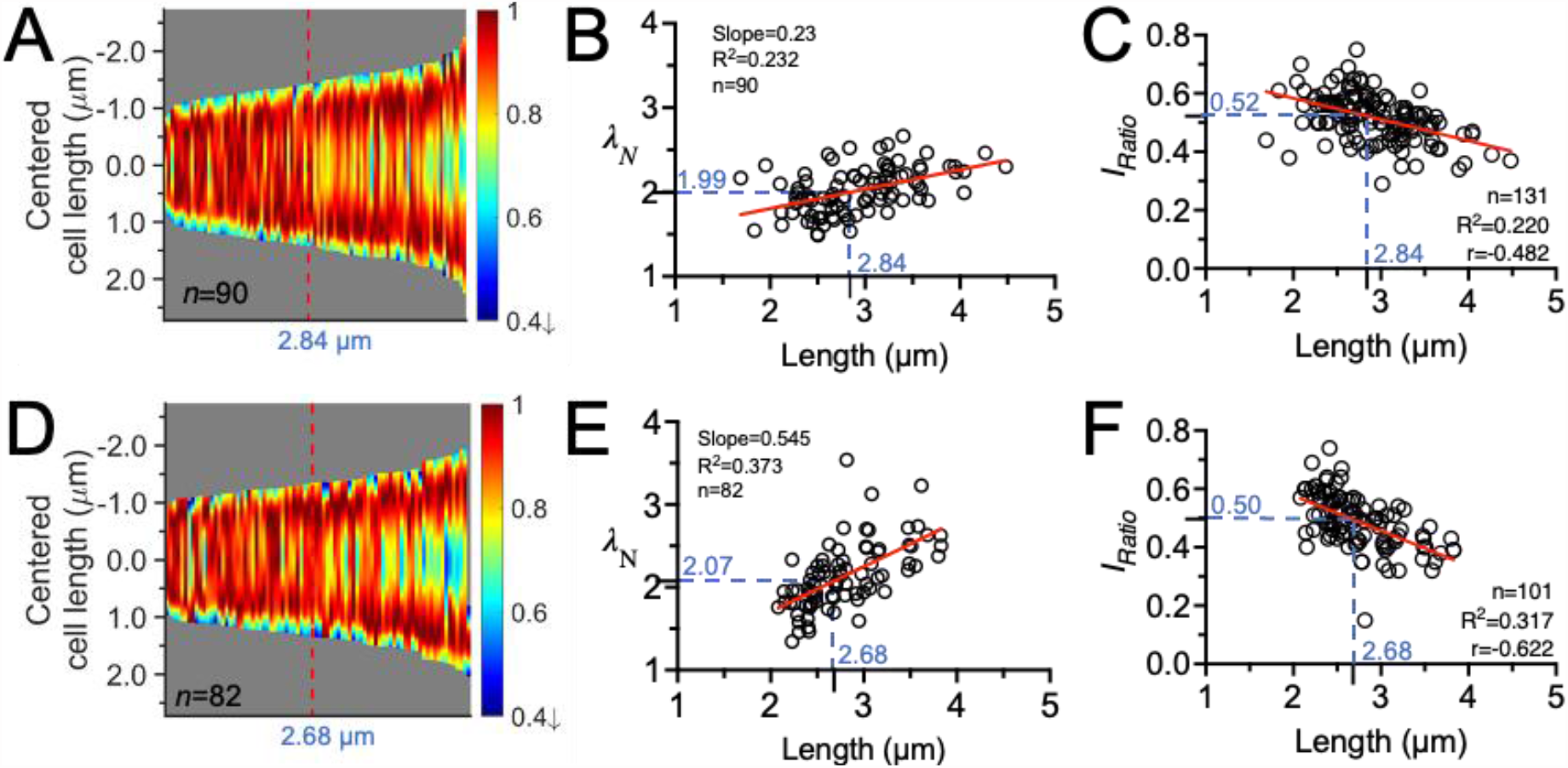
Characterization of MinD concentration gradients. Cells were grown in 0.4% (A-C) and 0% glucose (D-F). **A**,**D**, Demographs of the fitted intensity profiles. The red line in each graph indicates the median cell length in the population. **B**,**E**, Correlation plot of wave slope (*λ*_*N*_) versus cell length. **C**,**F**, Correlation plot of intensity ratio (*I*_*Ratio*_) versus cell length. In B, C, E and F, the red connecting lines were obtained by simple linear regression. The blue dashed lines represent the *λ*_*N*_ or *I*_*Ratio*_ values at the median cell length. n, population size; R^2^, goodness of fit; r, Spearman coefficient.

Next, the decay constant *λ*_*N*_ of the concentration gradient was calculated to numerically describe the wave slope of the MinD concentration gradient (Fig. S2). The *λ*_*N*_ value was estimated over the normalized cell length that became dimensionless in order to allow comparisons between cells of different lengths. As demonstrated in Fig. 3B, smaller *λ*_*N*_ (gentler slope) is found in shorter cells and larger *λ*_*N*_ (steeper slope) is found in longer cells, suggesting that as cells get longer, the wave slope towards the center of the cell becomes steeper. These observations led us further to hypothesize that a variable MinD concentration gradient could couple with a changing MinD concentration at the midcell to promote septum formation.

Therefore, the second numerical descriptor is the intensity ratio (*I*_*Ratio*_), defined as the ratio between the minimum and maximum fluorescence values determined in the intensity profile. The minimum intensity *I*_*min*_ was determined at the intersection of paired concentration gradients (Fig. S2), which was located in the vicinity of the midcell with very few exceptions. The maximum intensity *I*_*max*_ was approximately 14.9 ± 4.5% of the cell length from either pole. The decrease in fluorescence near the poles may be due to the smaller volume of the hemispherical poles (Fig. S3E). The *I*_*Ratio*_ value calculated according to *I*_*min*_ and *I*_*max*_ reflects the amount of MinD accumulation near the cell middle and can be used to predict the probability of septum formation. As shown in Fig. 3C, *I*_*Ratio*_ is inversely proportional to cell length, indicating a progressively greater difference in MinD concentration at poles and division sites in growing cells.

### Steeper concentration gradient in shorter cells under glucose starvation

Similarly, variable concentration gradients and a decrease in the midcell concentration of MinD were also observed in cells grown under glucose starvation (Fig. 3D-F), even when the oscillation period was slow (Fig. 1G). Quantitatively, *λ*_*N*_ values were larger and *I*_*Ratio*_ values decreased faster in cells cultured with 0% glucose (Fig. 3E, F), suggesting that a steeper concentration gradient under glucose starvation may favor division at shorter cell lengths. However, the *I*_*Ratio*_ values at the median cell length showed no significant difference between nutrient shifts (Fig. 3C, F), indicating that the *I*_*Ratio*_ value at division fell within a similar range, disregard of division at shorter cell lengths under glucose starvation or longer cell lengths under normal condition.

It should be noted that fluctuations in intracellular ATP concentrations due to glucose downshifts are unlikely to affect the amount of ATP required to maintain MinD oscillations. While the cellular concentration of ATP under different conditions was reported as 1-5 mM (Lasko & Wang, 1996; Mathis & Brown, 1976; Schneider & Gourse, 2004; Soini *et al*, 2005), the rate of ATP consumption was reported as∼0.9 μmole/μmole MinD/sec (∼30 nmole/mg MinD/min) (Shih *et al*, 2011). Thus, the abundance of ATP in cells greatly exceeds the ATP consumed by MinD even under glucose starvation.

### Simulating dynamic concentration gradient of MinD in growing cells

A one-dimensional mathematical model based on Fange and Frey (Fange & Elf, 2006; Wu *et al*., 2015) was employed to reveal physical insights variable concentration gradients as cells elongate. This model describes the major reactions underlying the oscillating Min system (Fig. 4A), including attachment of cytosolic MinD-ATP to the membrane, recruitment of MinD-ATP and MinE by membrane-bound MinD, dissociation of the MinDE complex from the membrane, and nucleotide exchange with ATP in cytosolic MinD (Materials and Methods and Supplemental Information). Each reaction is associated with a combination of different parameters that determine the reaction conditions in the simulation (Table S4, Excel Table 1), including the concentrations of MinD and MinE experimentally measured in this study, the diffusion coefficients of Meacci et al. (2005, 2006) (Meacci & Kruse, 2005; Meacci *et al*, 2006), previously simulated dissociation rate constant *k*_*de*_ for MinDE complexes (Fange & Elf, 2006; Wu *et al*., 2015), and the searched parameter set consisted of four rate constants, *k*_*dD*_, *k*_*dE*_, *k*_*D*_, and *k*_*ADP→ATP*_, which were scanned with random numbers to probe the general behaviors of the system.

**Fig. 4.**
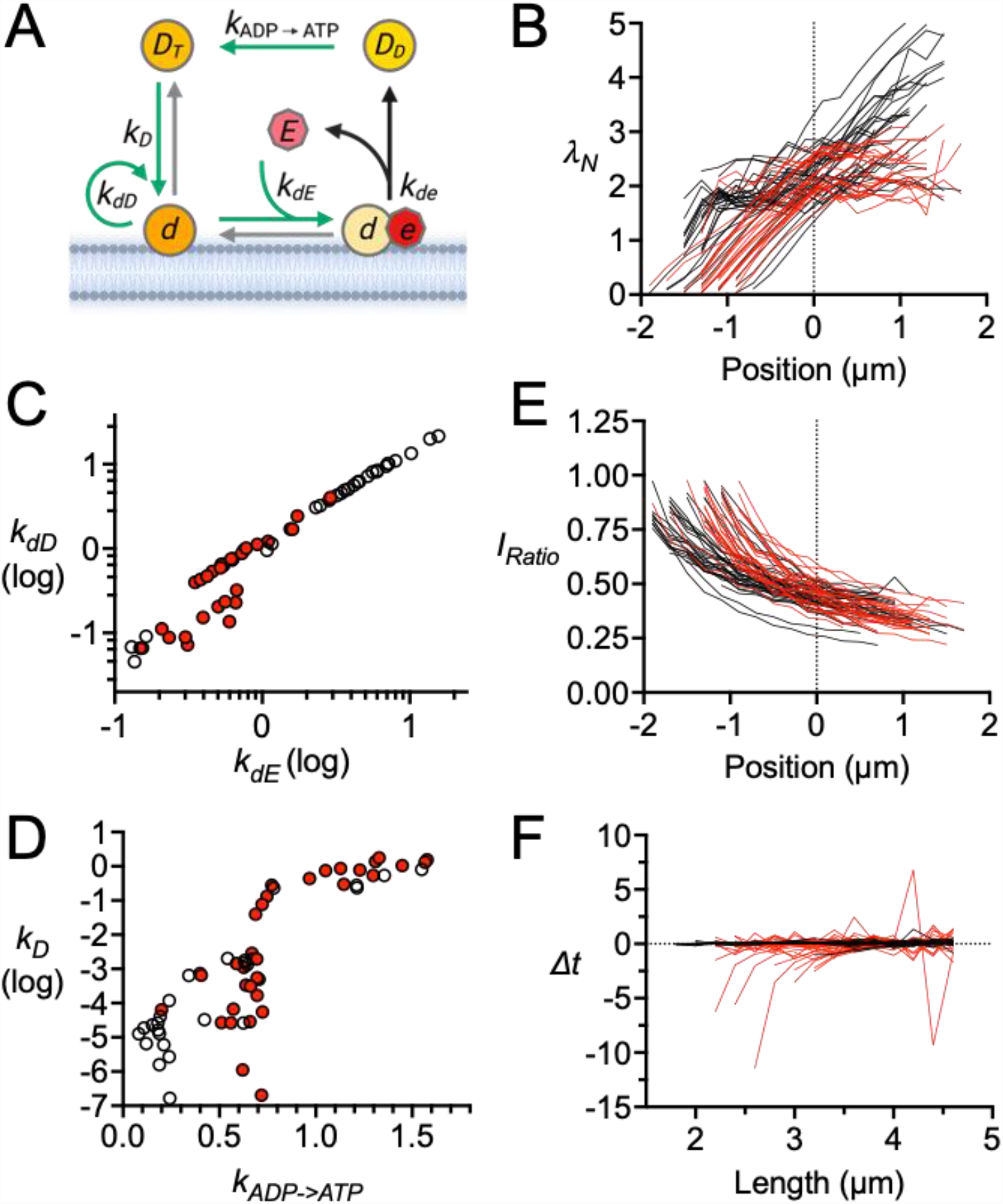
*in silico* characterization of MinD concentration gradients. **A**, Illustration of the molecular interactions between MinD and MinE and the corresponding reaction rate constants in the mathematical model. While the rate constant *k*_*de*_ is fixed in the simulation (black arrow), other four rate constants (*k*_*dD*_, *k*_*dE*,_ *k*_*D*_, and *k*_*ADP→ATP*_; green arrows) are variable values. Gray arrows are reactions that were not considered. **B**, Plot of *λ*_*N*_ versus position. The biphasic lines are aligned at the transition length (*L*_*t*_) that resets to the zero position. **C**, Correlation plot between the rate constants *k*_*dD*_ and *k*_*dE*_. **D**, Correlation plot between the rate constants *k*_*D*_ and *k*_*ADP→ATP*_. **E**, Plot of *I*_*Ratio*_ versus position. The biphasic lines are aligned at *L*_*t*_ as in B. **F**, The oscillation period of virtue cells hardly changed with increasing length. In B-F, uniphasic (n=27) and biphasic (n=33) modes are represented in black and red, respectively.

A linear stability analysis was performed on the model, requiring the most divergent linear solution to oscillate in time and space as it approaches steady state (Supplemental Information, Fig. S6). Of the 5000 parameter sets, 638 passed the linear stability criterion and exhibited spatiotemporal oscillations. Of these, 174 parameter sets yielded *λ*_*N*_ and *I*_*Ratio*_ values that matched experimental observations in at least 4 different cell lengths (Fig. S6A). The vast majority of these parameter sets show *λ*_*N*_ as a function of length that are further divided into two groups according to the fitted curve (Fig. S6B). The uniphasic group shows the increment *λ*_*N*_ of the fitted linear function (Fig. 4B black line, S7A), while the biphasic group shows the increment *λ*_*N*_ of the first phase and the approximation of the second phase (Fig. 4B, S7B). In all cases, the incremental phase required a mean relative error (<0.12) and an upper slope limit of 2 μm^-1^ (Fig. S6A).

### Variable concentration gradients and their coupling to division site placement

The data processing described above yielded 60 parameter sets, 27 for the uniphasic group and 33 for the biphasic group (Fig. 4B-F, S7), showing different combinations of four rate constants (Excel Table 1). In general, *k*_*dD*_ and *k*_*dE*_, representing the rates at which membrane-bound MinD recruits MinD-ATP and MinE, are tightly coupled to balance the amount of MinD and MinE on the membrane (Fig. 4C). When comparing the rate constants in the uniphasic and biphasic groups, *k*_*dD*_ and *k*_*dE*_ in the biphasic mode are significantly lower than those in the uniphasic mode (Fig. 4C), while the nucleotide exchange (*k*_*ADP→ATP*_) in MinD in the biphasic mode is higher than that in the uniphasic mode (Fig. 4D). Taken together, the results suggest that establishment of a biphasic pattern requires slower recruitment of MinD and MinE to the membrane and faster nucleotide exchange with ATP to recharge MinD.

Subsequent analyses involving the numerical descriptors *λ*_*N*_ and *I*_*Ratio*_ allow for a better understanding of the coupling between variable concentration gradients and the division site placement (Fig. 4B, E, S6, S7). We found that important features are preserved regardless of uniphasic or biphasic modes, including the length dependence of *λ*_*N*_ and *I*_*Ratio*_, and increasing *λ*_*N*_ in combination with decreasing *I*_*Ratio*_ that can facilitate the placement of the FtsZ ring. The conclusions from this analysis are consistent with our *in vivo* measurements, which occur in cells under normal conditions (Fig. 3B) and are exacerbated in carbon-starved cells (Fig. 3E).

### The oscillation period hardly changes with the cell length

As shown in Fig. 4F and S8, the analysis also supports maintenance of the oscillation period as cells elongate. Interestingly, parameter sets with smaller *k*_*dD*_ and *k*_*dE*_ more frequently produced longer oscillation periods in both uniphasic and biphasic groups. From the period*−*length plots (Fig. S8), most oscillation periods appear to be insensitive to changes in cell length (Fig. 4F), and this property is completely independent of the parameters tested in the current study.

This finding is supported by a linear stability analysis of the mathematical model, where the most divergent solution close to the steady state oscillates with frequency, which varies slowly with the cell length *L*, as predicted by the imaginary part of the corresponding eigenvalue (Fig. S9, Supplemental Information). Thus, maintenance of the oscillatory period during cell elongation is a universal property found in both living bacteria and numerical simulations.

## Discussion

Understanding oscillatory dynamics in the cellular environment is necessary to bridge the gap created by differences in spatial scale and organization *in vivo* and *in vitro*. This study revealed the plasticity of the MinD concentration gradient in growing *E. coli* cells (Fig. 5). A length-dependent dynamic concentration gradient combined with a gradual decrease in the concentration of MinD at the midcell (Fig. 5) may act in synchrony with MinC to allow placement of the FtsZ ring. We also found that the variable concentration gradient is accompanied with relatively stable oscillation period and increasing velocity (Fig. 1E, F), while the accuracy of division site placement is ensured (Fig. S5). These characteristics of MinD oscillation are maintained under carbon stress (Fig. 1G). In addition, there may be a default speed of MinD oscillation in newborn cells, as indicated by the slower speed after cell division (Fig. S4).

**Fig. 5.**
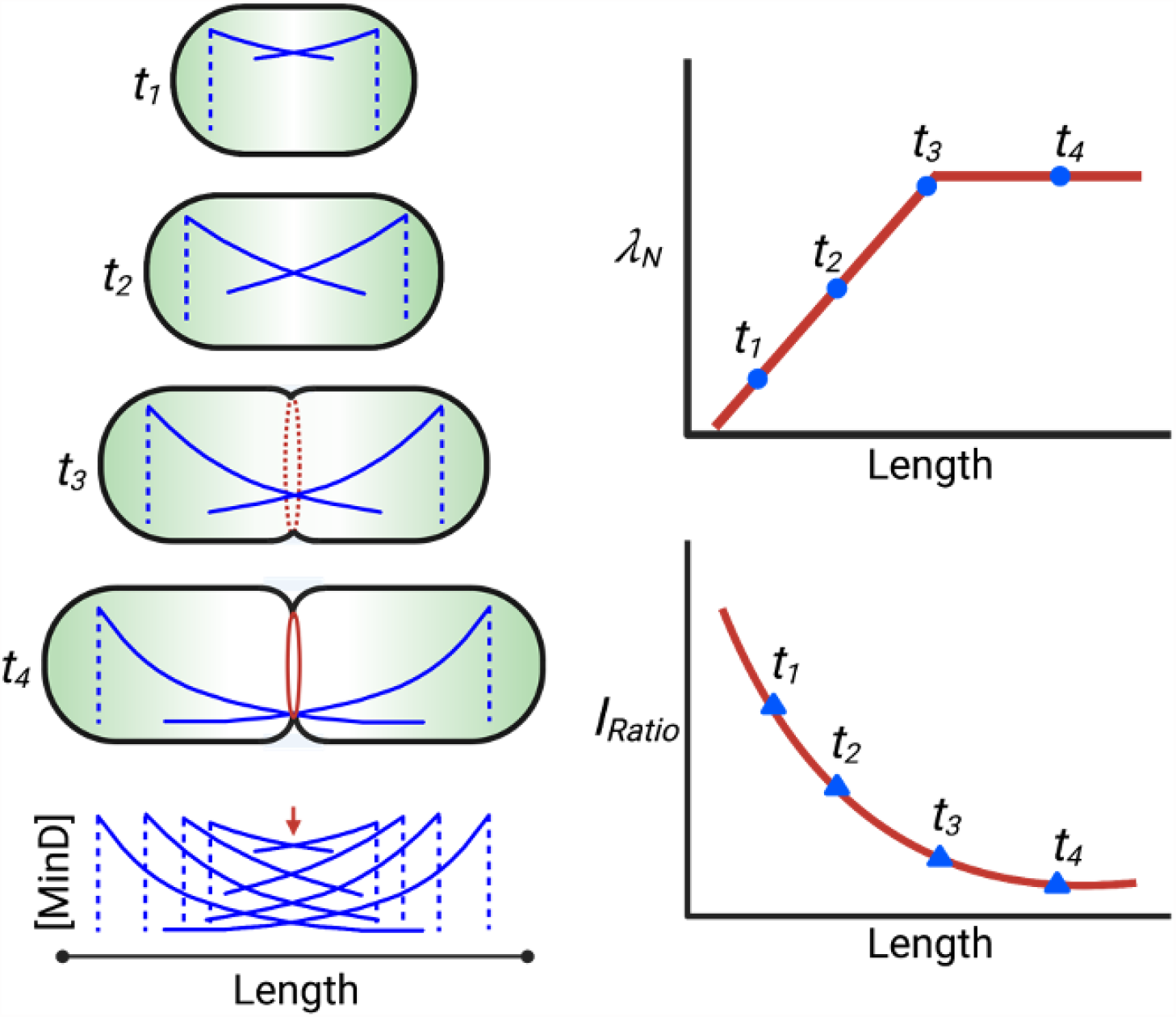
Schematic representation of the tunable MinD concentration gradient during the cell cycle. Green shading: MinD concentration; blue line: concentration distribution; red dashed and solid lines: FtsZ ring; red arrow: position of lowest MinD concentration.

Concentrations of MinD and MinE are key determinants of oscillation patterns of the Min system (Lutkenhaus, 2007; Denk *et al*., 2018; Mizuuchi & Vecchiarelli, 2018). Our observations of maintaining the oscillatory period and increasing velocity are consistent with the *in vitro* reconstitution study of Vecchiarelli et al. (Vecchiarelli *et al*., 2016), where period maintenance is explained by increasing the concentration of MinE to promote the release of MinD from the membrane. To gain further insight into variable concentration gradients, we measured the cellular concentrations of MinD and MinE (Fig. S1) and calculated the number of MinD molecules throughout the cell cycle (Fig. 2B, C). Although there are technical difficulties in imaging MinE at the cellular level in parallel, knowing changes in MinD concentration and number of molecules in growing cells is sufficient to understand variable concentration gradients. Our measurements showed that the cellular concentration of MinD per unit area remains constant as cells grow (Fig. 2B), suggesting the concentration gradient of MinD (rather than the concentration itself) is more critical for the change in gradient shape.

In this study, the concentration gradient of MinD is characterized by numerical descriptors *λ*_*N*_ and *I*_*Ratio*_, representing the wave slope and the intensity ratio between the minimum and maximum values in the gradient, respectively (Fig. S2). While *λ*_*N*_ showed a progressively increasing trend with increasing length (Fig. 3B), *I*_*Ratio*_ showed a gradually decreasing trend with increasing length (Fig. 3C). The same trend was found in cells cultured under carbon-rich and starved conditions, both of which support changing MinD concentration gradients as an intrinsic feature of the Min system.

A kinetic model of the Min oscillation aids in understanding the underlying mechanisms of period maintenance and variable concentration gradients (Fig. 4A, Supplemental Information). The simulation results supported that the oscillation period was largely independent of length (Fig. S8), as the oscillation frequency derived from the linear stability analysis showed only small variations at different lengths covering the range of normal cell lengths (Fig. S9). For the shape descriptors of the concentration gradient, *λ*_*N*_ and *I*_*Ratio*_, simulation results showed similar trends to the experimental observations. Thus, the mathematical model yields an adequate characterization of the observed kinetics and successfully reinforces the experimental observations involving variable concentration gradients and constant oscillation periods.

Based on the identification of the transition point *L*_*t*_ between the incremental slope to the plateau in the *λ*_*N*_*−*Length plot, the parameter sets obtained from the simulation were divided into uniphasic and biphasic groups, each group exhibiting different preference in combinations of the four parameters (Fig. 4C, D, S6). Interestingly, the biphasic character of the MinD concentration gradient is associated with slower recruitment of MinD and MinE by the membrane-bound MinD (smaller *k*_*dD*_ and *k*_*dE*_ rate constants), but faster nucleotide exchange to recharge MinD with ATP (larger *k*_*ADP→ATP*_ rate constant).

Furthermore, since the movement of spatial boundaries in the numerical model represents cell growth *in vivo*, one might wonder whether the decay curve of the concentration gradient initially emerging from the end of the cell changes with cell length. If so, the decay curve of the MinD concentration gradient would be a plateau where the decay curve remains constant, which is called the scaling behavior of the concentration gradient. Interestingly, this scaling behavior occurs during the second phase of the biphasic mode, during which approximate *λ*_*N*_ values, indicative of scaling behavior, are observed (Fig. 4C). This result indicates the concentration gradient can be scaled proportionally to cell length in longer cells, which can maintain MinD concentration at low levels suitable for FtsZ ring formation. Such biphasic nature of the MinD concentration gradient may be advantageous, as a faster decline (steeper *λ*_*N*_) may favor cells coping with irregular oscillations, especially in newborn cells. As shown in Fig. 1 and a previous study (Juarez & Margolin, 2010), regular interpolar oscillations of the Min system occur more frequently in longer cells but not in newly dividied cells. The flexibility between the two modes may be a strategy to tune the MinD concentration gradient in order to regulate cell division under different growth conditions. Nonetheless, unlike the perfect scaling model of morphogen gradients that provide relative positional signals in embryo and tissue development (Capek & Muller, 2019; Inomata, 2017), the scaling behavior of MinD concentration gradients is to maintain concentrations appropriate for FtsZ ring placement.

In conclusion, this work reveals the plasticity of the MinD concentration gradient as an intrinsic property of the Min system during pole-to-pole oscillations throughout cell growth. This plasticity arises from spatial differences in molecular interactions between MinD and MinE, as demonstrated by using experimentally measured cellular concentrations of MinD and MinE to search for combinations of rate constants for different molecular interactions in a mathematical model. We also show that this variable concentration gradient, coupled with the clearance of MinD at the midcell, prepares cells for division as they grow, further distinguishing *in vivo* and *in vitro* observations.

## Materials and methods

### Strains and plasmids

Genotypic descriptions of strains and plasmids are provided in Table S1. The primer sequences are provided in Table S2.

The *E. coli* strains MC1000 and DH5α were used for general cloning purposes. The strain BL21(DE3)/pLysS was used for protein production. Strain FW1541 (*ΔminD minE::sfgfp-minD minE::frt aph frt*) (Wu *et al*., 2015) was derived from the laboratory strain W3110 (*F-λ-IN(rrnD-rrnE)1 rph-1*).

### Growth conditions

Each bacterial culture was grown from a single colony in minimal medium containing M9 salts (Difco™), 0.25% casamino acids, 2 mM MgSO_4_, and 0.1 mM CaCl_2_ and supplemented with 0.416% glucose at 30°. The overnight culture was used to inoculate fresh medium of the same kind to an OD_600 nm_ value of approximately 0.05 and then allowed to grow at 30°C until the OD_600 nm_ reached between 0.3 and 0.4. Cells were washed twice with M9 salt solution and resuspended in 30 μL of minimal medium supplemented with the desired concentration of glucose. Three microliters of cell suspension were then spotted on a 2% agarose pad prepared on a glass slide. The agarose pad was prepared with the desired concentration of glucose to investigate the concentration effect on the sfGFP-MinD oscillation (0.416% or 0%). After the agarose pad was sealed under a coverslip, and the slide was placed on a preheated stage at 30°C installed on the microscope and allowed to stabilize for 10 min before image acquisition.

Antibiotics were used at the following concentrations when plasmids were handled: 50 μg/mL ampicillin, 50 μg/mL kanamycin, and 34 μg/mL chloramphenicol. Anhydrotetracycline (0.1 μM) was used for induction of the P*LtetO-1* promoter.

### Microscopy

Our microscopy system included an Olympus IX81 inverted microscope (Olympus, Tokyo, Japan) equipped with a CCD camera (ORCA-R2, Hamamatsu, Japan), an objective lens (UPlanFLN 100x, NA 1.30, Olympus, Tokyo, Japan), and filter sets for sfGFP (ET-Narrow Band EGFP filter set, cat. 49020, Chroma Technology Corporation, VT, USA) and mScarlet-I (Semrock LF561-A-OMF, IDEX Health & Science, LLC Center of Excellence, Rochester, New York, USA). Images were acquired using Xcellence Pro software (Olympus Corp., Tokyo, Japan) with an X-Cite® 120 Metal Halide lamp (Excelitas Technologies Corp., Ontario, Canada) or cellSens Dimension software (Olympus Corp., Tokyo, Japan) with an X-Cite^®^ TURBO multiwavelength LED illumination system (Excelitas Technologies Corp., Ontario, Canada).

Time-lapse images of sfGFP-MinD were acquired with a 12-sec interval for 10 min or before the fluorescence diminished. The images were analyzed using the MicrobeJ plugin v5.11j (Ducret *et al*, 2016) installed in NIH ImageJ (Rasband, W.S., ImageJ, U.S. National Institutes of Health, Bethesda, Maryland, USA, http://rsb.info.nih.gov/ij/). In brief, the time-lapse images were converted into a demograph compiled from the one-dimensional fluorescence intensity profiles generated by the projection of fluorescence onto the medial axes of the same cell. This measurement obtained the ensemble fluorescence of sfGFP-MinD in the membrane and in the cytosol on an axial position. The oscillation period was measured from the demograph. Only oscillation cycles with clear starting and ending points were used in the subsequent statistical analyses.

### Statistical analyses

Statistical analyses of the experimental data were performed in GraphPad Prism (Version 8.3, GraphPad Software, Inc., USA). Spearman correlation method was used to calculate the correlation between the paired data with a 95% confidence interval to obtain the correlation coefficient (r) and two-tailed probability value (P). For comparing datasets obtained from FW1541 under different culture conditions, the nonparametric Kruskal-Wallis test (significance level *α*=0.05) and Dunn’s multiple comparisons test were used. The number of data points and any data exclusion for each experiment are explained in the figures, the figure legends, or the tables.

### Image processing

Image processing was performed using MATLAB R2018b (The MathWorks, Inc., USA). The intensity *I* was collected as 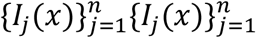 with a time stamp 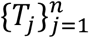 associated with the individual intensity datum and the position *x* ∈ *L*_*j*_, where *L*_*j*_ is a set to collect the observation positions of *x* and is normalized by setting that *(L*_*j*_*) =*0 and *(L*_*j*_*) =* 1 for each *j*. Further data processing shifts all *I*_*j*_(x), to have its minimum value 0 over *x*.

As the intensity has been observed to decay over time due to photobleaching, we scale the intensity function by a factor 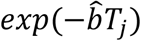 where 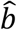 is the fitted value of the decay rate *b* in the model to describe the decay behavior due to photobleaching, resulting in *f(t) = a · exp(−bt)*. We obtain 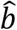 by taking the regression of *f(t)* values at time stamps 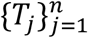 on the total intensity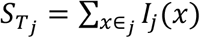 with *j =* 1, *…, n*. Thus, we have 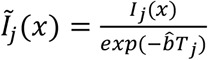 for each *j =* 1, *…, n* after data processing.

To quantify changes of 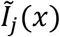 over *j* (the time stamp index), that is expressed as 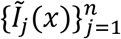, we introduce an index 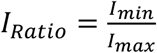, where *I*_*min*_ and *I*_*max*_ are the minimum and maximum values of the summarized curve for 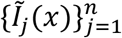. We split the data in two groups according to their left or right position, by applying the k-mean analysis on the slope estimations at position 0.2 and 0.8 and excluding those with small slopes within the first quartile range. We then apply Gaussian kernel smoothing technique to these two position groups, respectively, and output the corresponding summary curves 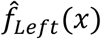 and 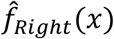 as well as 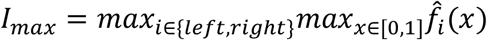.

To obtain *I*_*min*_, defined as the intensity at the intersection of the left and right curves, we employ the exponential decay model to have a robust estimation. We apply the equation 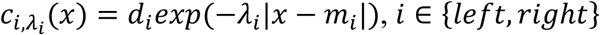 to fit the two groups of data by taking 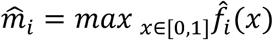. We measure the overall decay rate by the weighted average *λ*^*′*^ of *λ*_*left*_ and *λ*_*right*_ for *ĉ*_*Left,λ*_*′* and *ĉ*_*Right,λ*_*′*, in which the weights are the sizes for these two groups. That is 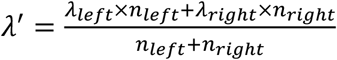. Thus, *I*_*min*_ is the intensity of the intersection of *ĉ*_*Left,λ*_*′* and *ĉ*_*Right,λ*_*′*. Then, 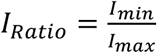 is calculated.

### Determination of the cellular concentration of sfGFP-MinD by fluorescence imaging

The cellular concentration of sfGFP-MinD was studied by culturing FW1541 in M9 minimal medium supplemented with 0.416% glucose, and the image acquisition and processing methods were performed as described earlier. Snapshots of both phase-contrast and fluorescence channels were acquired every 15 min for 5 hours. The sum intensity in individual cells and the intensity per unit area (μm^2^) were corrected as described below and plotted against time to understand the changes in fluorescence intensity during the cell cycle (Fig 2A and B).

The time-lapse fluorescence sequences were corrected for photobleaching. The intensity 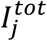 measured from a cell is defined as a function of *k*,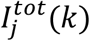, where *k* refers to rounds of photobleaching (*k ≥ 0*) caused by light exposure applied to the *j*^*th*^ cell (*j =* 1, *…, n*). *I*_*j*_*(k)* is a decreasing function of *k*, since exposure to light can cause irreversible damage to the fluorophores, resulting in a gradual decrease in the fluorescence intensity. The normalized intensity, 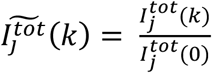, is obtained by dividing 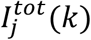 by its maximum 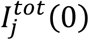. Given 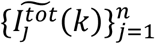, we employed a biexponential curve model 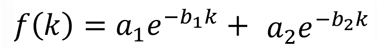 to obtain the parameters 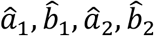 to build the reference curve 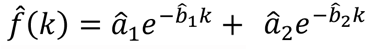. The image data obtained from the time-lapse sequence, *C*_*j*_*(k)*, were corrected for photobleaching. The intensity of the *j*^*th*^ cell after photobleaching correction was *Ĉ*_*j*_*(k)*, where 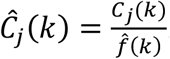. A linear regression model was applied to the data obtained from individual cells using the following formula: 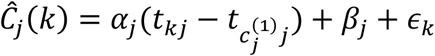, where 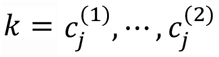. Here, 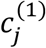 is the first image data point of a cell acquired after the first cell division, and 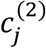 is the last image data point of the same cell acquired before the next cell division. Therefore, 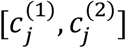 is the number of image frames or data points between two rounds of cell division. *t*_*kj*_ is the time when the *j*^th^ cell undergoes the *k*^th^ round of light exposure. Then, the overall trend for a population of cells (*n*) was estimated using 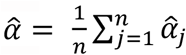 and 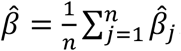.

### Modeling

A mathematical model was developed based upon rate constants of the following reactions to examine the MinD concentration gradient at different cell lengths (Fig. 4A, S6, Supplemental Materials and Methods).

1. Attachment of cytosolic MinD-ATP (MinD.ATP_c_) to the membrane and becoming a membrane-bound form (MinD.ATP_m_):

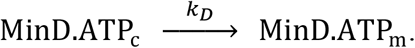
2. Recruitment of MinD.ATP_c_ by MinD.ATP_m_:

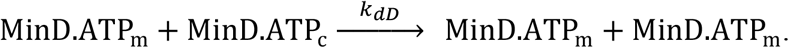
3. Recruitment of cytosolic MinE (MinE_c_) by MinD.ATP_m_ and forming a membrane-bound MinDE complex (MinDE_m_):

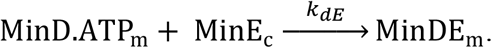
4. Dissociation of MinDE_m_ from the membrane through the ATP hydrolysis in MinD and becoming MinD.ADP_c_ and MinE_c_:

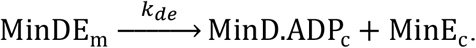
5. Nucleotide exchange in MinD.ADP_c_ and becoming MinD.ATP_c_:

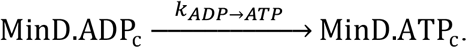

The mass action kinetics for the concentrations *cx*, with *x*=*DD, DT, E, d* and *de*, denoting MinD.ADP_c_, MinD.ATP_c_, MinE_c_, MinD.ATP_m_ and MinDE_m_, respectively, with their diffusion is:

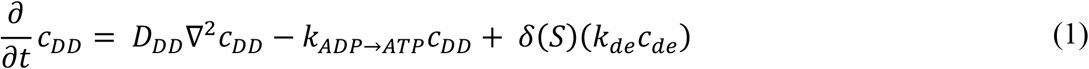

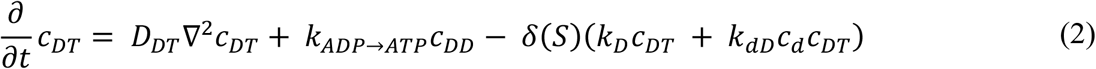

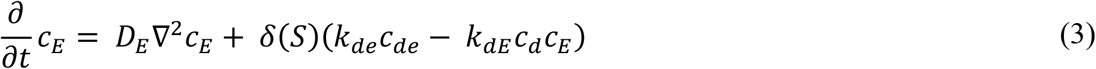

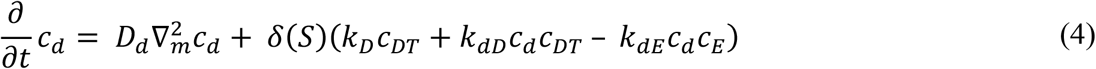

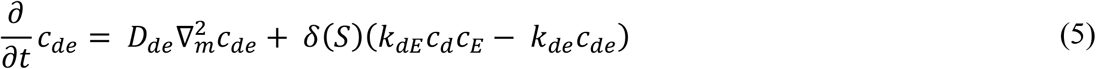

Here, we followed Wu et al, 2015 (Wu *et al*., 2015) and use *δ(S)* to denote the membrane reactions. Eqs. 1-5 were solved numerically with a simple finite difference scheme in a one-dimensional model, with grid size 0.2 μm. The dynamic simulation was performed initially under a length of 3 μm to search for parameters, and subsequently scanned through a length range from 1.6 to 4.6 μm. The numbers of the MinD and MinE molecules were set to 2205 (1.95 μM) and 1580 (1.4 μM) (Fig. S1), respectively, at the experimental median cell length 2.84 μm (Table 1), which was changeable proportionally with the length in the simulation. The rate constant *k*_*de*_ was set at 0.33 (1/sec) (Wu *et al*., 2015), since we found that the general time scale of the dynamics is sensitive to *k*_*de*_. Other reaction rates needed in the model were searched with random samplings. The diffusion coefficients for MinD and MinE in the cytosol are 16 and 10 μm^2^/sec and those in the membrane are 0.2 μm^2^/sec (Meacci *et al*., 2006). The other four rate constants, *k*_*D*_ (1/sec), *k*_*dD*_ (μm/sec), *k*_*dE*_ (μm/sec) and *k*_*ADP→ATP*_ (1/sec), were randomly searched. Initial tests were performed with a linear stability analysis and promising parameter sets, consisting of the 4 rate constants that show both Turning and Hopf instability, are selected for further tests in the actual dynamics.

The numerical fitting procedures for obtaining *λ*_*N*_ and *I*_*Ratio*_, and the detailed condition for the selection of the final parameter sets in this work are described in Supplemental Information and Fig S6.

## Acknowledgments

We thank Cees Dekker and Harold Erickson for providing plasmids and strains, Ester Malau for assistance in protein purification, and Jian Liu, Min Wu, Chien-Jung Lo, and Todd Lowary for comments and discussion. This work was funded by the National Science and Technology Council, Taiwan, through grants to YLS (NSTC 110-2311-B001-011, 108-2311-B001-012, 107-2311-B001-023, 106-2311-B001-009), IPT (NSTC 106-2118-M-001-001-MY2), and CPH (NSTC 111-2123-M-001-003), and by the Grand Challenge Seed Project program of Academia Sinica through a grant to IPT (AS-GCS-108-08). YLS and CPH also acknowledge intramural support from Academia Sinica, Taiwan.

## Conflict of Interests

The authors declare that they have no conflict of interest.

## Materials availability

The plasmids, genetically engineered *E. coli* strains, and reagents generated in this study are available from the lead contact with a completed materials transfer agreement. There are restrictions on the availability of the anti-MinD and anti-MinE antisera because the purification procedures result in small quantities.

## Data and code availability

All image datasets and their measurements are available for review upon request. The computer scripts for image processing and modeling the Min protein oscillation are available from the following links, https://drive.google.com/drive/folders/1O-nlT4AlJen2UHRh2GcrrfqQuc0ykgsH?usp=share_link and https://zenodo.org/record/8122833, respectively.

## Figures and figure legends

**Excel Table 1.** Sampled reaction rate constants output from simulated MinD oscillations. Data relate to **Fig. 4C, D**.

| Bi-phasic (33) | $k_{de}$ | $k_D$ | $k_{AD}$ | $k_{AE}$ | $k_{ADP \rightarrow ATP}$ |
| --- | --- | --- | --- | --- | --- |
| 0011-bi | 0.33 | 1.372618 | 0.22947 | 0.660982 | 1.30993 |
| 0098-bi | 0.33 | 0.861912 | 0.204223 | 0.503166 | 1.128331 |
| 0453-bi | 0.33 | 0.775109 | 0.087987 | 0.233212 | 1.228976 |
| 0770-bi | 0.33 | 2.68E-05 | 0.951019 | 0.747384 | 0.510115 |
| 0793-bi | 0.33 | 2.67E-05 | 2.420804 | 1.728459 | 0.559098 |
| 0875-bi | 0.33 | 5.54E-05 | 0.655772 | 0.541896 | 0.72303 |
| 1064-bi | 0.33 | 0.132598 | 1.706945 | 1.552193 | 0.746604 |
| 1093-bi | 0.33 | 0.076854 | 1.2126 | 1.097543 | 0.721703 |
| 1453-bi | 0.33 | 2.87E-05 | 0.397976 | 0.350176 | 0.658506 |
| 1487-bi | 0.33 | 0.001104 | 0.429291 | 0.380999 | 0.626589 |
| 1706-bi | 0.33 | 0.000335 | 0.63728 | 0.540802 | 0.637377 |
| 1728-bi | 0.33 | 0.27787 | 1.692037 | 1.603505 | 0.771501 |
| 1886-bi | 0.33 | 0.000496 | 0.630113 | 0.544675 | 0.705961 |
| 2274-bi | 0.33 | 0.438963 | 0.321798 | 0.670235 | 0.96797 |
| 2298-bi | 0.33 | 6.69E-05 | 0.758359 | 0.632945 | 0.572274 |
| 2426-bi | 0.33 | 0.002862 | 0.874409 | 0.726926 | 0.668525 |
| 2519-bi | 0.33 | 0.746826 | 0.234334 | 0.559727 | 1.050336 |
| 2540-bi | 0.33 | 0.001964 | 0.535698 | 0.458316 | 0.673822 |
| 2920-bi | 0.33 | 0.001442 | 0.643974 | 0.526596 | 0.588281 |
| 2923-bi | 0.33 | 6.33E-05 | 3.734038 | 2.832188 | 0.199423 |
| 2933-bi | 0.33 | 0.297809 | 0.111897 | 0.208722 | 1.145093 |
| 2962-bi | 0.33 | 0.001947 | 0.470664 | 0.422626 | 0.693835 |
| 3118-bi | 0.33 | 1.11E-06 | 1.012406 | 0.778097 | 0.620324 |
| 3161-bi | 0.33 | 2.03E-07 | 1.121398 | 0.925868 | 0.718716 |
| 3553-bi | 0.33 | 0.540242 | 0.066196 | 0.153655 | 1.297851 |
| 3766-bi | 0.33 | 0.000551 | 0.73135 | 0.622029 | 0.695408 |
| 3789-bi | 0.33 | 1.540903 | 0.072308 | 0.311168 | 1.577067 |
| 4153-bi | 0.33 | 1.301561 | 0.089211 | 0.301252 | 1.566651 |
| 4294-bi | 0.33 | 1.048451 | 0.152455 | 0.397277 | 1.448715 |
| 4299-bi | 0.33 | 0.000168 | 0.752665 | 0.616483 | 0.695039 |
| 4762-bi | 0.33 | 0.000311 | 0.592094 | 0.521269 | 0.661545 |
| 4873-bi | 0.33 | 1.758964 | 0.136355 | 0.599268 | 1.32812 |
| 4963-bi | 0.33 | 0.277481 | 0.091081 | 0.162993 | 1.211469 |
| Uni-phasic (27) | $k_{de}$ | $k_D$ | $k_{AD}$ | $k_{AE}$ | $k_{ADP \rightarrow ATP}$ |
| 0236-uni | 0.33 | 0.000118 | 8.167366 | 5.983729 | 0.239421 |
| 0300-uni | 0.33 | 2.31E-05 | 7.189122 | 5.189392 | 0.152226 |
| 0407-uni | 0.33 | 3.28E-05 | 6.088879 | 4.426729 | 0.422806 |
| 0618-uni | 0.33 | 2.61E-05 | 6.226307 | 4.54416 | 0.180521 |
| 0843-uni | 0.33 | 0.000734 | 3.076703 | 2.311699 | 0.399708 |
| 0934-uni | 0.33 | 2.69E-06 | 19.97589 | 13.72316 | 0.239307 |
| 0975-uni | 0.33 | 0.541192 | 0.065868 | 0.148591 | 1.355239 |
| 0984-uni | 0.33 | 0.231351 | 0.954146 | 1.075795 | 0.781914 |
| 1097-uni | 0.33 | 0.002027 | 3.226804 | 2.467922 | 0.542915 |
| 1376-uni | 0.33 | 1.28E-05 | 4.90557 | 3.619357 | 0.18832 |
| 1552-uni | 0.33 | 1.27E-05 | 13.42093 | 10.23017 | 0.081435 |
| 1574-uni | 0.33 | 0.039466 | 1.138586 | 1.160495 | 0.687555 |
| 2073-uni | 0.33 | 0.001515 | 5.633228 | 4.17418 | 0.632831 |
| 2606-uni | 0.33 | 1.87E-05 | 10.96032 | 7.994681 | 0.108311 |
| 2637-uni | 0.33 | 1.64E-07 | 8.067043 | 5.5818 | 0.242594 |
| 2980-uni | 0.33 | 4.01E-05 | 3.918919 | 2.894558 | 0.19204 |
| 3056-uni | 0.33 | 1.62E-05 | 8.433492 | 6.039562 | 0.181223 |
| 3276-uni | 0.33 | 0.000646 | 4.259758 | 3.207386 | 0.341947 |
| 3479-uni | 0.33 | 0.235627 | 0.06798 | 0.130143 | 1.213931 |
| 3663-uni | 0.33 | 1.59E-06 | 10.13371 | 7.123297 | 0.188387 |
| 3702-uni | 0.33 | 2.62E-05 | 4.544762 | 3.442072 | 0.624503 |
| 3798-uni | 0.33 | 0.001335 | 5.030857 | 3.817781 | 0.644481 |
| 3923-uni | 0.33 | 6.04E-06 | 10.12645 | 7.192824 | 0.211795 |
| 4193-uni | 0.33 | 0.807925 | 0.045504 | 0.136096 | 1.549716 |
| 4334-uni | 0.33 | 6.53E-06 | 21.50707 | 15.66315 | 0.120675 |
| 4345-uni | 0.33 | 0.001782 | 9.473695 | 6.92035 | 0.631853 |
| 4854-uni | 0.33 | 0.000652 | 4.020668 | 2.89876 | 0.405425 |

## Notes

### Competing Interest Statement

The authors have declared no competing interest.

