## Supplemental Information for "Growth-dependent concentration gradient of the oscillating Min system in *Escherichia coli*"

### Supplemental Materials and Methods

*Strain and plasmid construction*

The strain SOT88 (W3110 *ΔminC minD minE*) was constructed following the standard procedures of the λ Red recombination method (Datsenko & Wanner, 2000). The plasmid pKD3 was used as the template to generate the recombinant DNA fragment using primers that carried homologous sequences 50 bp upstream of the ATG start codon of *minC* and the terminal 50 bp of *minE* as well as the primer sequences on pKD3. The PCR product was transformed into W3110/pKD46 competent cells and screened for recombinants. The procedure resulted in strain SOT87 (W3110 *ΔminC minD minE, cat*), which was cured for pKD46 followed by removal of the *cat* cassette to generate the strain SOT88. The flanking DNA fragments of *minCDE* were PCR-amplified for sequencing to confirm the deletion of the three genes.

The plasmid pSOT370 ($P_{LtetO\text{-}1}$::*ftsA-mScarlet-I*) was generated using the Gibson assembly method (Gibson Assembly® Master Mix, New England Biolabs) to ligate the DNA fragments of *ftsA* that were PCR-amplified from the genomic DNA, *mScarlet-I,* which was amplified from pEB2-mScarlet-I (Balleza *et al*, 2018), and pMLB1113-$P_{LtetO\text{-}1}$, which was the large fragment of pSOT329 digested with SmaI and HindIII ($P_{LtetO\text{-}1}$::$ftsZ^{G55}$-*mKO2*-$ftsZ^{Q56}$). The plasmid pSOT329 ($P_{LtetO\text{-}1}$::$ftsZ^{G55}$-*mKO2*-$ftsZ^{Q56}$) was modified from pSOT295 ($P_{lac}$::$ftsZ^{G55}$-*mKO2*-$ftsZ^{Q56}$) by replacing the *lac* promoter with the *LtetO-1* promoter from pdCas9 bacteria (Qi *et al*, 2013). pSOT295 was created by replacing *mCerulean* in pSOT294 ($P_{lac}$::$ftsZ^{G55}$-*mCerulean*-$ftsZ^{Q56}$) with *mKO2,* which was amplified from pSOT291 ($P_{lac}$::*ftsZ-mKO2*). pSOT291 originated from pSOT157 ($P_{lac}$::*ftsZ-yfp)* after *yfp* was replaced with *mKO2* amplified from FW2454 (*hupA-mKO2*) (Wu *et al*, 2015a). pSOT157 was constructed by three-fragment ligation between the XbaI and HindIII fragments from pMLB1113 (de Boer *et al*, 1989), the XbaI-*ftsZ*-BamHI fragment that was amplified from the *E. coli* genome by PCR and subjected to restriction digestion, and the BamHI-*yfp*-HindIII fragment from pYLS67 (Shih *et al*, 2002). The *mCerulean* gene was amplified by PCR from pJSB-*FtsZ*$^{G55}$-*mCerulean-FtsZ*$^{Q56}$ (Moore *et al*, 2017).

For the construction of pSOT279 ($P_{lac}$::*his*$_{6x}$-*sfgfp-minD*), *sfgfp-minD* was amplified from pBVS4 ($P_{lac}$::*sfgfp-minD minE*) (Wu *et al*, 2015b), and the *his*$_{6x}$ tag and restriction sites NheI and BamHI were introduced using primers. The resulting PCR product was restriction digested and ligated into pET21a.

*Growth condition of studying ftsA localization*

To study FtsA localization, FW1541/pSOT370 cells were grown from a single colony in M9 minimal medium supplemented with 0.416% glucose and 50 µg/ml ampicillin at 30°C. FtsA was studied in place of FtsZ because of the abnormal morphology caused by the overexpression of *ftsZ* or by engineering *ftsZ* on the chromosome. The overnight culture was used to spike the fresh medium and allowed to grow until mid-log phase. Cells were spun down, washed, and diluted to $OD_{600\ nm}$ ~0.2 in the same medium. Then, 0.1 µM anhydrotetracycline (aTc) was added to induce the expression of *ftsA-mScarlet-I* from pSOT370. The culture was incubated for 2.5 hours at 30°C before being spun down and resuspended in 30 µL of medium. Three microliters of cell suspension were spotted on an agarose pad containing M9 medium with 0.416% glucose and 0.1 µM aTc for imaging. The strain was imaged for FtsA localization before acquiring the MinD oscillation.

*Characterization and purification of the anti-MinD antiserum*

The fusion protein Trx'-His$_{6x}$-MinD was purified as described in Hsieh et al., 2010 (Hsieh *et al*, 2010) and used to raise rabbit polyclonal antiserum by LTK Biolaboratories (Taoyuan County, Taiwan). Then, 50 mL of the final bleed was collected from the immunized rabbit. This crude antiserum was purified against purified $His_{6x}$-MinD using the blotting method. $His_{6x}$-MinD was overexpressed from strains BL21(DE3)/pLysS/pSOT6 ($P_{lac}$*::his*$_{6x}$*-minD*) and purified following the procedure described in Shih et al., 2019 (Shih *et al*, 2019). Thirty micrograms of $His_{6x}$-MinD were run on a 10% SDS-PAGE gel and transferred onto a PVDF membrane (0.45 µm, Amersham GE Healthcare Europe GmbH, Germany) using a TE 22 Mighty Small™ Transphor Tank Transfer Unit (Amersham GE Healthcare Biosciences Inc., USA). The membrane was stained with 0.5% Ponceau S to visualize the protein for excision of the membrane band containing $His_{6x}$-MinD. The membrane band was washed three times with TBST (20 mM Tris-Cl, pH 7.4, 150 mM NaCl, 0.05% Tween 20) and blocked in 10 mL of 5% bovine serum albumin (BSA) prepared in TBST. The membrane band was incubated with 0.5 mL of crude antiserum diluted in 9.5 mL of TBST at 4°C with gentle shaking overnight. The band was washed three times with 10 mL of TBST before stripping off the antibodies from the blot using 1 mL of 0.2 M glycine/HCl (pH 2.5) and immediately neutralized with 1 mL of 1 M Tris-Cl, pH 9.0. The purified antiserum was validated against total cell lysates of FW1541 and W3110 as well as against $His_{6x}$-sfGFP-MinD and $His_{6x}$-MinD purified by Western blotting, as shown in Fig S1A.

*Characterization and purification of the anti-MinE antiserum*

Overexpression and purification of the fusion protein MinE-$His_{6x}$ from the strain BL21(DE3)/pLysS/pSOT13 ($P_{lac}$*::minE-his*$_{6x}$) were performed following previously described procedures (Shih *et al.*, 2019). The purified MinE-$His_{6x}$ was used to raise the rabbit polyclonal antiserum as described for the anti-MinD antiserum. The crude antiserum was purified against the cell lysate of strain SOT88 to remove nonspecific contaminants, followed by purification against MinE-$His_{6x}$. In brief, 100 µg of SOT88 cell lysate and 30 µg of MinE-$His_{6x}$ were separated on 15% Tris-Tricine gels and transferred independently onto PVDF membranes. The membrane was stained with 0.5% Ponceau S for visualization and excision of the membrane band containing SOT88 cell lysate or MinE-His$_{6x}$. Both membrane bands were treated as described for anti-MinD antiserum.

Crude antiserum (0.5 mL) diluted in 9.5 mL of TBST was first incubated with the membrane band of SOT88 cell lysate with gentle shaking at 4°C for 8 h. After removal of the first membrane band that absorbed nonspecific contaminants in the antiserum, the solution was incubated with the second membrane band containing MinE-His$_{6x}$ with gentle shaking at 4°C overnight. This membrane band was washed before stripping off the anti-MinE antibodies as described above. The purified antiserum was validated against the total cell lysates of W3110, FW1541 and SOT88 cells as well as MinE-His$_{6x}$ purified by Western blotting, as shown in Fig S1B. The antiserum was divided into aliquots and stored in 10% (v/v) glycerol at -20°C.

*Determination of cellular concentrations of sfGFP-MinD and MinD by Western blotting*

One liter of exponentially growing cells was spun down at 6,000 ×*g* for 20 min at 4°C and washed 3 times with 30 mL of 1x PBS (137 mM NaCl, 2.7 mM KCl, 10 mM $Na_2HPO_4$, 1.8 mM $KH_2PO_4$, pH 7.4). The cell pellet was resuspended in 20 mL of lysis buffer [50 mM Tris-Cl pH 7.5, 500 mM NaCl, 100 µg/mL lysozyme, 200 µg/mL DNase I, 1 protease inhibitor tablet (cOmplete™, EDTA-free Protease Inhibitor Cocktail, Roche, Basel, Switzerland)]. This cell suspension was incubated on ice for 30 min before disruption using a NanoLyzer N2 (Gogene Corporation, Hsinchu, Taiwan) with two passages at 15,000 psi. The clear lysate was recovered after centrifugation at 15,000 ×*g* for 10 min at 4°C and quantified using the Bio-Rad Protein Assay Kit (Bio-Rad Laboratories, Inc., CA, USA).

His$_{6x}$-MinD and His$_{6x}$-sfGFP-MinD were overexpressed from strains BL21(DE3)/pLysS/pSOT6 (Shih *et al.*, 2019) and BL21(DE3)/pLysS/pSOT279 ($P_{lac}$::*his*$_{6x}$-

*sfgfp-minD*), respectively, and purified following the procedure described in Shih et al., 2019 (Shih *et al.*, 2019). The purified fusion proteins of MinD were used to generate concentration standards for quantifying the cellular concentrations of MinD and sfGFP-MinD in strains W3110 and FW1541.

Forty micrograms of clear lysate along with a serial dilution of the purified $His_{6x}$-sfGFP-MinD or $His_{6x}$- MinD were separated on a 10% SDS-PAGE gel, followed by blotting onto a PVDF membrane. Three batches of cell lysates and three repeats of each batch were analyzed for statistical purposes. The membrane was blocked with 5% BSA in TBST (50 mM Tris-Cl, pH 7.4, 150 mM NaCl, 0.1% Tween 20) for an hour at room temperature before incubation with the purified anti-MinD antiserum at a 1:100 dilution at 4°C overnight. The blot was washed 3 times with TBST and incubated with 10,000-fold diluted horseradish peroxidase-conjugated anti-rabbit IgG antibody (Amersham ECL rabbit IgG, HRP-linked whole Ab from donkey, Cytiva–Global Life Sciences Solutions, Marlborough, USA) at room temperature for one hour. After washing 5 times with TBST, the blots were treated with ECL reagents (Amersham Biosciences$^{TM}$ ECL Select$^{TM}$ Western Blotting Detection Reagent, GE Healthcare). The bioluminescence signal was detected using an ImageQuant LAS-4000 system (GE Healthcare Life Sciences).

The band intensity of the blots was quantified by NIH ImageJ. The measurements from serial dilutions of $His_{6x}$-sfGFP-MinD and $His_{6x}$-MinD were used to generate a calibration curve to estimate the amounts of sfGFP-MinD and MinD in the lysate by linear regression. The cellular concentrations (M) of sfGFP-MinD and MinD were calculated by dividing the amount of protein in grams by the molecular weight (sfGFP-MinD: 56,817 Da; MinD: 29,614 Da), the number of cells in 40 μg of cell lysate, and the single-cell volume (Fig S3E). The number of protein molecules per cell was obtained by multiplying the result by Avogadro's number.

The cellular concentrations of sfGFP-MinD and MinD were quantified from 3 independent exponential cultures of FW1541 and W3110, respectively. The dry weight of three independent samples was measured from cells collected from 50 mL cultures that were freeze-dried overnight at 10 mTorr (UNISS Freeze Dryer FDM-20, Taiwan Green Version Technology Ltd., Taiwan). The wet weight (g/mL) was estimated from the dry weight based on an assumption that water accounts for 75% of the cell weight (Bionumbers ID 105482; https://bionumbers.hms.harvard.edu/bionumber.aspx?id=105482). The estimation of the single-cell weight is shown in Table S3.

*Determination of cellular concentrations of MinE by Western blotting*

Sixty micrograms of clear lysate along with a serial dilution of the purified MinE-His$_{6x}$ were separated on a 15% Tris-Tricine gel, followed by the procedures described above for MinD. The molecular weight of MinE (10,235 Da) was used in the calculation.

*Linear stability analysis of the numerical model*

As described in the main text, we performed the linear stability analysis (Cross, 2006) on the five-chemical reaction-diffusion model in order to search for the parameters. This analysis is based on the Hopf bifurcation theories that predict a fixed point near oscillation. With diffusion, the uniform solution becomes unstable and an oscillation pattern appears with a specific finite wave length, a condition we aimed to follow in finding suitable parameters.

For simplicity, we reduced the diffusion-reaction problem to one dimension, in which $x$ denotes the position along the cell's long axis. Thus, Eqs. 1−5 as described in the main text can be rewritten as the vector form:

$$\frac{\partial}{\partial t}\mathbf{u} = \mathbf{D}\frac{\partial^2}{\partial x^2}\mathbf{u} + \mathbf{f}(\mathbf{u}) \quad \text{(S1)}$$

with the diffusion matrix $\mathbf{D}$ defined as:

$$\mathbf{D} = \begin{pmatrix} D_{DD} & 0 & 0 & 0 & 0 \\ 0 & D_{DT} & 0 & 0 & 0 \\ 0 & 0 & D_E & 0 & 0 \\ 0 & 0 & 0 & D_d & 0 \\ 0 & 0 & 0 & 0 & D_{de} \end{pmatrix}.$$

and a vector function $\mathbf{f}(\mathbf{u})$ denoting the reaction expressions as shown in Eqs. 1−5.

For oscillation dynamics to occur, it is generally assumed that there is a uniform fixed point near the steady state, which denoted as $\mathbf{u}^*$, with $\mathbf{f}(\mathbf{u}^*) = 0$. After Taylor expansion, we kept only up to the linear terms near the steady state $\mathbf{u}^*$ and rewrote Eq. S1 as a set of linear equations with a constant, A:

$$\frac{\partial}{\partial t}\delta\mathbf{u} = \mathbf{D}\frac{\partial^2}{\partial x^2}\delta\mathbf{u} + \mathbf{A}\,\delta\mathbf{u} \quad \text{(S2)}$$

where $\mathbf{A} = \partial\mathbf{f}/\,\partial\mathbf{u}$, a 5 by 5 Jacobian matrix evaluated at $\mathbf{u}^*$. With Eq. S2 being a linear equation for $\delta\mathbf{u}$, a standard procedure is plugged in the following trial solution:

$$\delta\mathbf{u} = \delta\mathbf{u}_q e^{\sigma_q t} e^{iqx}, \quad \text{(S3)}$$

where $\delta\mathbf{u}_q$ is an arbitrary initial condition expressed as a vector. Substituting the expression in Eq. S3 with $\delta\mathbf{u}$ in Eq. S2, we can obtain the following eigenvalue problem:

$$\mathbf{A}_q\delta\mathbf{u}_q = \sigma_q\delta\mathbf{u}_q, \quad \text{(S4)}$$

with the Jacobian $\mathbf{A}_q$ defined as:

$$\mathbf{A}_q = \mathbf{A} - \mathbf{D}q^2 =$$

$$\begin{pmatrix} -k^{\mathrm{ADP\to ATP}} - D_{DD}q^2 & 0 & 0 & 0 & k_{de} \\ k^{\mathrm{ADP\to ATP}} & -k_D - k_{dD}c_d^* - D_{DT}q^2 & 0 & -k_{dD}c_{DT}^* & 0 \\ 0 & 0 & -k_{dE}c_d^* - D_E q^2 & -k_{dE}c_E^* & k_{de} \\ 0 & k_D + k_{dD}c_d^* & -k_{dE}c_d^* & k_{dD}c_{DT}^* - k_{dE}c_E^* - D_d q^2 & 0 \\ 0 & 0 & k_{dE}c_d^* & k_{dE}c_E^* & -k_{de} - D_{de}q^2 \end{pmatrix}$$

(S5)

Namely, the eigenvalue ($\sigma_q$) of $\mathbf{A}_q$ is solved as a function of $q$, which describes the stability of such a solution. When projected to its eigenvector, a positive real part of $\sigma_q$ indicates an increased deviation of $\delta\mathbf{u}_q$ with time, and a negative real part of $\sigma_q$ indicates a decreased deviation of $\delta\mathbf{u}_q$. Meanwhile, a complex value of $\delta\mathbf{u}_q$ is a solution for oscillation in time.

*Linear stability analysis for the oscillation frequency*

A typical result of linear stability analysis is demonstrated in Fig S9. When the wave vector $q$ is varied and the eigenvalues of the linearized dynamics are near a fixed point, both the real and the imaginary parts of the eigenvalues are marked with the largest real part. As a result, the nonzero frequency is predicted to dominate over a range of the cell lengths. Although the actual nonlinear dynamics may differ significantly from the linearized prediction, we used the nonlinear analysis to test whether the wave shape, or the concentration gradient, would scale with length. The analysis can reveal a scaling behavior that the preferred wavelength of the system matches the cell length, or the non-scaling behavior with an unmatching wavelength that could arise from modulated reaction rates and diffusion of the molecules. Nevertheless, the actual oscillation frequency and the range of cell length that permits oscillation are quite similar to those predicted by the linear analysis, represented by eigenvalues, for most of the parameter sets tested (Fig S9).

*Screening of the rate constants that generate MinD oscillation*

The initial 5000 parameter sets, as described in the main text, were first screened for a divergent oscillating solution in time and space under the linear stability analysis. In linear dynamics, the eigenvalue with the largest real part prevails at a long time, and thus an oscillation in time is predicted. Such an eigenvalue, which is sometimes called a Hopf instability, has a non-zero imaginary part and a positive real part. When it happens for an oscillatory trial solution in space, $e^{iqx}$, a Turing instability is predicted. To facilitate the search of parameters showing features similar to the Min oscillation, the linear stability analysis along with the constraints of both Hopf unstable (in time) and Turing unstable (in space) conditions were employed. Namely, we calculated the steady state using its corresponding Jacobian matrix (Eq. S5), with $q = \pi/L$. Judging from the eigenvalues, the parameter sets from those with the most divergent component or namely with a positive real part and a nonzero imaginary part, were kept for full numerical simulation using Eqs. 1−5 as in the main text. Here, the most divergent solution in the linear approximation describes MinD oscillation between two cell ends with a wavelength of $2L$, where $L$ is the cell length under a least-oscillating condition that satisfies the zero-flux boundary condition. Under this condition, the wavevector $q$ is set to $\pi/L$ for these tests.

In the initial search, the cell length was set as 3 μm and the kinetic parameters, including the dissociation constant $k_{de}$, the diffusion coefficients $D_D$, $D_d$, and $D_{de}$, and concentrations of MinD and MinE, were fixed as listed in Table S4. Here, conversion of the protein concentrations between cytosolic and membrane locations became unnecessary due to reducing the reaction-diffusion model to a one-dimensional counterpart. Thus, the concentration in one-dimension, $c_x^*$, is equivalent to the molecule number divided by length (1/μm). When simulating under different cell-length regimes, the number of MinD and MinE molecules were scaled up proportionally with a fixed concentration ratio of MinD to MinE (Wu *et al.*, 2015b). The equations Eqs. 1−5 were propagated with a simple finite difference scheme, with a time step set as $3.587 \times 10^{-5}$ sec. To achieve a good numerical precision, we kept the concentration change under 5% with a spatial grid size of 0.2 μm. The four reaction rate constants, including $k_D$ (1/sec), $k_{dD}$ (μm/sec), $k_{dE}$ (μm/sec), and $k_{ADP \to ATP}$ (1/sec), were randomly searched as $10^N$, with $N$ being a random number dawn from a normal distribution with a zero mean and the standard deviation as 3.0.

*Analysis of the simulation results: period, $\lambda_N$ and $I_{Ratio}$*

To ensure stable oscillation, we ignored the first 40 seconds of each selected cases in the subsequent analysis. The intensity, $I(x, t)$, was defined as the concentration of $\mathrm{MinD.ATP_m}$ and $\mathrm{MinDE_m}$, was normalized to the maximal value in both time and space. The oscillation period was identified as the averaged time difference between two consecutive maximal *I* in the same grid from 40 sec to 80 sec. Within a grid, only oscillations with the amplitude change larger than 5% were accepted. The maximal *I* values identified from all time points were fitted with an exponential decay function to obtain the spatial profile:

$$I(x) = a\, e^{-\lambda_N x} + \mathrm{c}, \tag{S6}$$

Eq. S6 was fitted from its maximal (1) to minimal values in a cell, with *a* and *c* are fitting constants, and $x$ is the relative position that goes from 0 to 1 (Fig. S6B). This allowed calculation of $\lambda_N$. In some shorter cells, the oscillation was observed only in time but not in space, where same amount membrane-bound MinD were obtained across the cell length. In these cases, $\lambda_N$ was set to zero.

As the maximal *I* value ($I_{max}$) appears alternatively in either left and right poles of a cell, we overlapped two such profiles, and recorded the minimal value ($I_{min}$) from the merged profile was the simulated $I_{Ratio}$ (Fig. S6C, D).

### Supplemental tables

**Table S1**. List of strains and plasmids.

| Strain | Genotype | Source |
|---|---|---|
| DH5α | *ΔlacZ ΔM15 Δ(lacZYA-argF) U169 recA1 endA1 hsdR17(rK-mK+) supE44 thi-1 gyrA96 relA1* | (Taylor *et al*, 1993) |
| MC1000 | *araD139 Δ(araABC-leu)7679 galU galK Δ(lac)X74 rpsL thi* | (Casadaban & Cohen, 1980) |
| BL21(DE3)/pLysS | *str. B* $F^{-}$ *ompT gal dcm lon hsdSB*($r_B^-$ $m_B^-$) *λ(DE3 [lacI lacUV5-*$T7_{p07}$ *ind1 sam7 nin5]) [malB*$^{+}$*]*$_{K\text{-}12}$*(λS)* pLysS *[*$T7_{p20}$ *ori*$_{p15A}$*] cat* | (Studier & Moffatt, 1986) |
| W3110 | *F*$^{-}$ *λ*$^{-}$ *IN(rrnD-rrnE)1 rph-1* | (Bachmann, 1972) |
| FW1541 | W3110, *ΔminD minE::sfgfp-minD minE <> kan frt* | (Wu *et al.*, 2015b) |
| FW2454 | W3110, *hupA-mKO2::aph <> frt* | (Wu *et al.*, 2015a) |
| SOT87 | W3110, *ΔminC minD minE <> cat frt* | This study |
| SOT88 | W3110, *ΔminC minD minE <> frt* | This study |
| **Plasmid** | | |
| pMLB1113 | *ColE1/pBR/pUC, bla* | (de Boer *et al.*, 1989) |
| pET21a | f1, pBR322, $P_{T7}$, *bla* | Novagen |
| pKD3 | *oriR6Kγ, bla, rgnB, cat, FRT* | (Datsenko & Wanner, 2000) |
| pKD46 | *oriR101, repA101(ts),* $P_{ara}$*-gam-bet-exo, bla, araC, [tL3]* | (Datsenko & Wanner, 2000) |
| pCP20 | *oripSC101(ts), repA101(ts), [cI857](λ)(ts), bla, cat, FLP* | (Datsenko & Wanner, 2000) |
| pBVS4 | pVBS3, $P_{lac}$*::sfgfp-minD minE frt, aph, bla* | (Wu *et al.*, 2015b) |
| pEB2-mScarlet-I | *oripSC101,* P*proC:: mScarlet-I, aph* | (Balleza *et al.*, 2018) |
| pdCas9-bacteria | p15A, $P_{LtetO\text{-}1}$*-dcas9, cat* | (Qi *et al.*, 2013) |
| pJSB-*ftsZ*$^{G55}$-*mCerulean* -*ftsZ*$^{G56}$ | pJSB, $P_{ara}$*::ftsZ*$^{G55}$*- mCerulean -ftsZ*$^{Q56}$*, cat* | (Moore et *al.*, 2017) |
| pYLS67 | pFX55, $P_{lac}$::*cfp-minD minE-yfp*, *bla* | (Hsieh *et al.*, 2010) |
| pFX55 | $P_{lac}$::*minC minD minE-yfp*, *bla* | (Shih *et al.*, 2002) |
| pSOT13 | pET21a, $P_{T7}$::*minE-his*$_{6x}$, *aph* | (Hsieh *et al.*, 2010) |
| pSOT157 | pMLB1113, $P_{lac}$*::ftsZ-yfp*, *bla* | This study |
| pSOT279 | pET21a, $P_{lac}$*::his*$_{6x}$*-sfgfp-minD, bla* | This study |
| pSOT291 | pMLB1113, $P_{lac}$*::ftsZ-mKO2, bla* | This study |

| | | |
|---|---|---|
| pSOT294 | pMLB1113, $P_{lac}$::*ftsZG55-mcerulean-Q56, bla* | This study |
| pSOT295 | pMLB1113, $P_{lac}$::*ftsZG55-mKO2-Q56, bla* | This study |
| pSOT329 | pMLB1113, $P_{LtetO-1}$::$ftsZ^{55}$-*mKO2*-$ftsZ^{56}$*, bla* | This study |
| pSOT370 | pMLB1113, $P_{LtetO-1}$::*ftsA-mScarlet-I, bla* | This study |

**Table S2**. List of primers. Underlined: primer sequence for the target gene; double underlined: primer sequence for the cloning vector; **bold face**: restriction site.

| Primer | Sequence | Source | Purpose |
|---|---|---|---|
| 910F | AACATCATCGCGCGCTGGCGATGATTAATAGCTAATTG AGTAAGGCCAGGGTGTAGGCTGGAGCTGCTTC | This study | SOT87 |
| 911R | CAAGGCAGAGATAACTCTGCCTTGAAGATAAATGCGCT TTTACAGCGGGCCATATGAATATCCTCCTTA | This study | SOT87 |
| 339F | AATT**CCCGGG**ATGTTTGAACCAATGGAACTTACC (XmaI) | This study | pSOT157 |
| 336R | CT**GGATCC**ATCAGCTTGCTTACGCAGGAAT (BamHI) | This study | pSOT157 |
| 439F | AA**GGATCC**ATCAGTAAAGGAG (BamHI) | This study | pSOT157 |
| 309R | AGTGCC**AAGCTT**ATTTGTAC (HindIII) | This study | pSOT157 |
| 279F | CTA**GCTAGC**ATGCACCACCACCACCACCACGGAGGAA TCCGTAAAGGCGAAGAGCTG (NheI) | This study | pSOT279 |
| 279R | CGC**GGATCC**TCATCCTCCGAACAAGCGTTTGAG (BamHI) | This study | pSOT279 |
| 1297F | GAT**GGATCC**GTGAGTGTGAT (BamHI) | This study | pSOT291 |
| 1298R | GCC**AAGCTT**AGGAATGAGCT (HindIII) | This study | pSOT291 |
| 1299F | GAT**GGATCC**GTGAGTGTGATTAAACCAGAGA (BamHI) | This study | pSOT294 |
| 1300R | GCC**AAGCTT**AGGAATGAGCTACTGCATCTT (HindIII) | This study | pSOT294 |
| 1304F | CGC**CCCGGG**ATGTTTGAACCAATGG (XmaI) | This study | pSOT294 |
| 1305R | GCG**AAGCTT**TTAATCAGCTTGCTTA (HindIII) | This study | pSOT294 |
| 1307F | TTGGAGGATCCACC**CTCGAG**GTGAGTGTGATTAAACC AGAAATGAAG (XhoI) | This study | pSOT295 |
| 1308R | GTCTGGGTGGATCC**CTCGAG**GGAATGAGCTACTGCAT CTTC (XhoI) | This study | pSOT295 |
| 1448F | GAGCGGATAACAATTTCACACAGGA | This study | pSOT329 |
| 1449R | ATCCGCTCATGAGACAATAACCCTG | This study | pSOT329 |
| 1450F | GTCTCATGAGCGGATACGTCTCATTTTCGC | This study | pSOT329 |
| 1451R | AATTGTTATCCGCTCCCATAGATCCTTTCT | This study | pSOT329 |
| 1563F | TGATTACGAATT**CCCGGG**ATGATCAAGGCGACGGAC (SmaI) | This study | pSOT370 |
| 1564R | AGAGCCGCCAGAGCCGCCTGAAAACTCTTTTCGCAGCC | This study | pSOT370 |
| 1565F | GGCTCTGGCGGCTCTATCAGTAAAGGAGAAGCTGTG | This study | pSOT370 |
| 1566R | CGGCCAGTGCC**AAGCTT**TTATTTGTATAGTTCATCC (HindIII) | This study | pSOT370 |

**Table S3**. Estimation of the single-cell weight (n=3).

| | **FW1541** | **W3110** |
|---|---|---|
| Harvesting $OD_{600nm}$ | 0.355−0.362 | 0.348−0.363 |
| Cell count (CFU/mL) | $7.96 \times 10^{8}$ | $7.71 \times 10^{8}$ |
| Dry weight (mg)/50 mL of culture | 5.6±0.72 | 5.2±0.64 |
| Single-cell weight (g/cell) | $5.65 \times 10^{-13}$ | $5.37 \times 10^{-13}$ |
| Cells/40 μg of lysate | $7.09 \times 10^{7}$ | $7.45 \times 10^{7}$ |

**Table S4**. Fixed parameter in the numerical simulation.

| Parameter | | Value (unit) | Source |
|---|---|---|---|
| Dissociation constant | $k_{de}$ | 0.33 (1/sec) | (Wu *et al.*, 2015b) |
| Diffusion coefficient | $D_D$ | 16 ($\mu m^2$/sec) | (Meacci *et al*, 2006) |
| | $D_d$ | 0.2 ($\mu m^2$/sec) | |
| | $D_{de}$ | 0.2 ($\mu m^2$/sec) | |
| MinD concentration | $c_{DD} + c_{DT} + c_d + c_{de}$ | 1.95 (μM) | This study |
| MinE concentration | $c_E + c_{de}$ | 1.4 (μM) | |

### Supplemental figures and figure legends

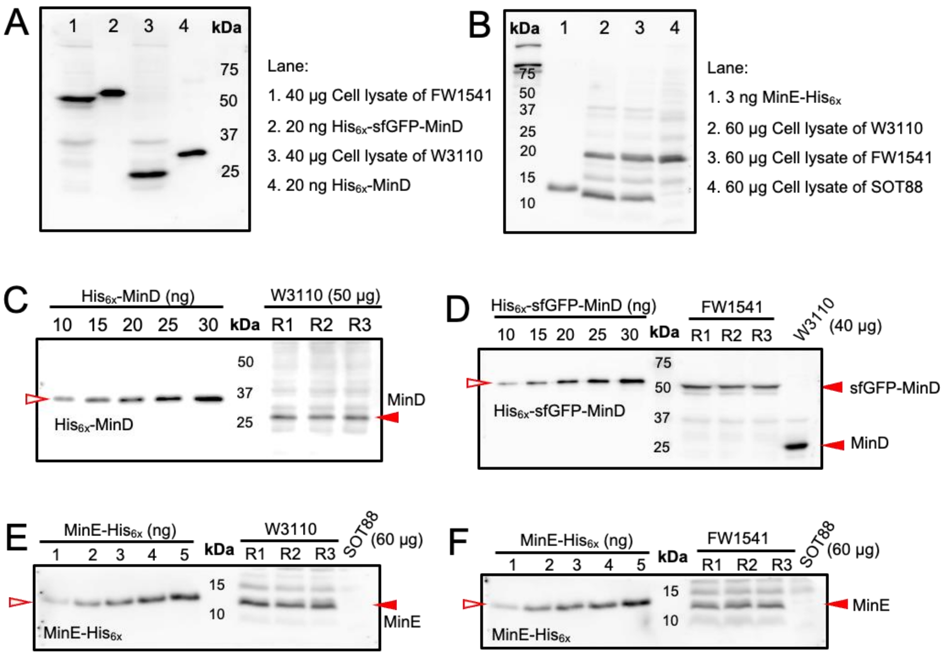


**Fig S1. Quantification of sfGFP-MinD, MinD and MinE.**

**A**, Validation of anti-MinD antiserum in detecting sfGFP-MinD and MinD in cell lysates alongside purified His$_{6x}$-sfGFP-MinD and His$_{6x}$-MinD. The blot-purified antiserum was used at a 1:100 dilution.

**B**, Validation of anti-MinE antiserum in detecting MinE in cell lysates alongside MinE-His$_{6x}$. The blot-purified antiserum was used at a 1:100 dilution.

**C**, **D**, An example of Western blots that was used to determine the concentrations of MinD in the cell lysates of the strains W3110 and FW1541, respectively.

**E**, **F**, An example of Western blots that was used to determine the concentrations of MinE in the cell lysates of the strains W3110 and FW1541, respectively. In C-F, serial dilutions of purified His$_{6x}$-MinD, His$_{6x}$-sfGFP-MinD, or MinE-His$_{6x}$ were applied to generate a calibration curve for interpolating the amount of MinD, sfGFP-MinD, or MinE in the sample by linear regression.

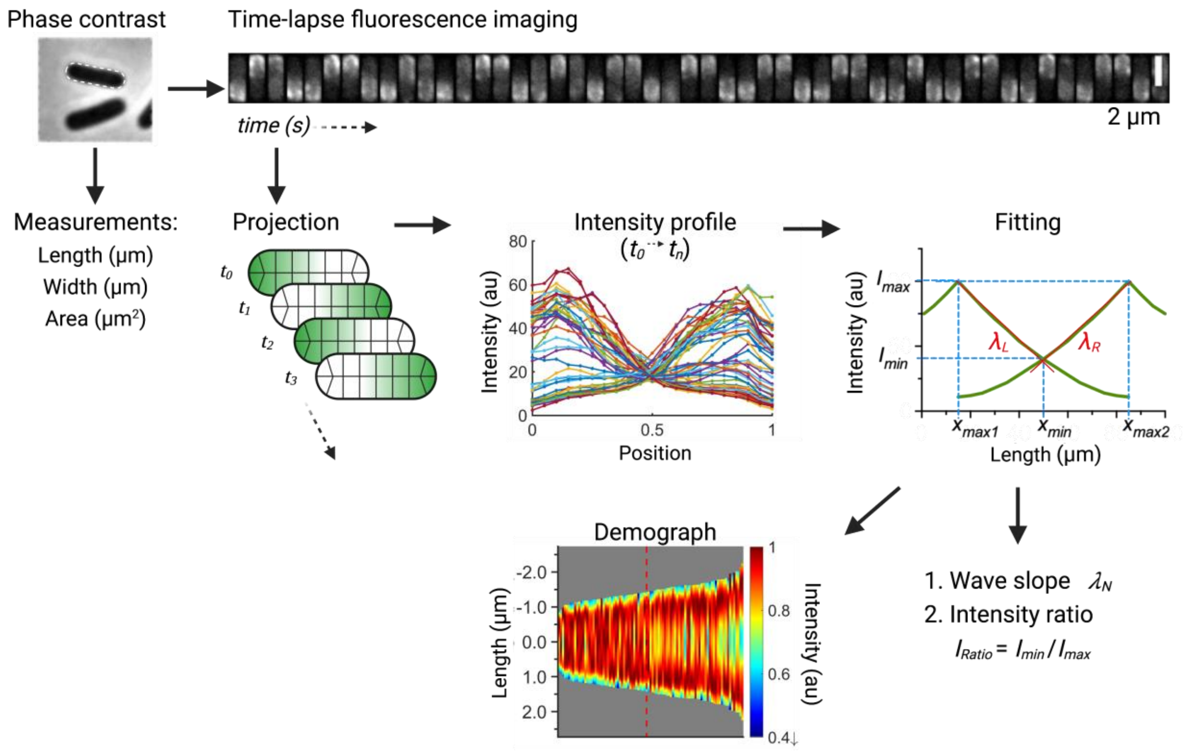


**Fig S2. Schematic diagram of image processing.**

Phase-contrast images were taken to measure cell size before acquiring time-lapse sequences of sfGFP-MinD oscillations. The fluorescent images were processed by projecting intensity values onto the median axis. The procedure was applied to all images throughout the time-lapse sequence. Then, the intensity at different axial positions was plotted to visualize fluorescence fluctuations, which indicate changes in cellular protein concentration. Next, a fitted curve was generated using the intensity profiles of the same cell in a time-lapse series that allowed measurements of the wave slope ($\lambda_N$) and the intensity ratio ($I_{Ratio}$). In addition, the fitted curves obtained from a population of cells were individually projected in one dimension, sorted by cell lengths, and compiled into a demograph.

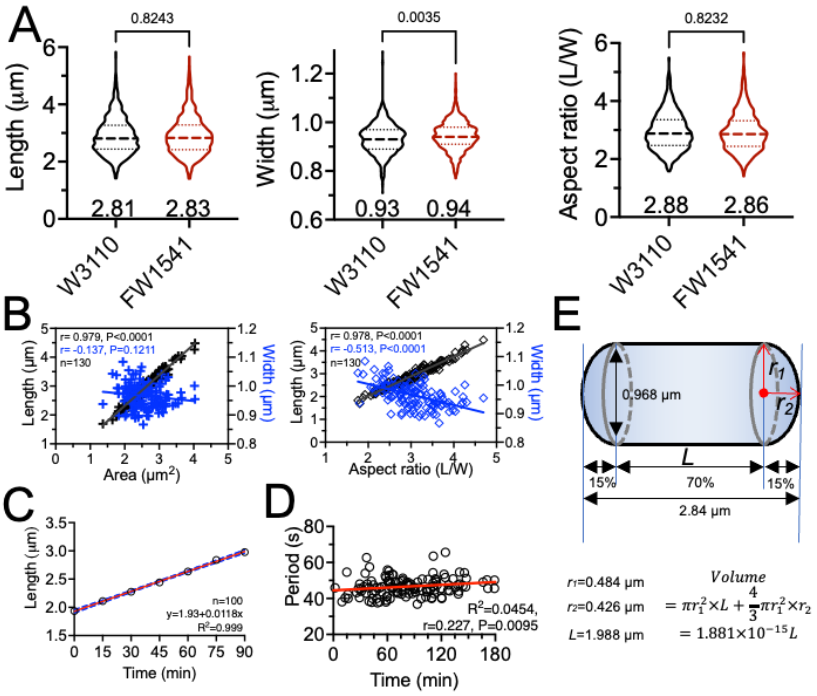


$$Volume = \pi r_1^2 \times L + \frac{4}{3}\pi r_1^2 \times r_2 = 1.881 \times 10^{-15} L$$

**Fig S3. Characterization of *E. coli* strains FW1541 and W3110 grown at 30°C in M9 medium supplemented with 0.416% glucose.**

**A**, Comparison of cell length, width, and aspect ratio. The median and interquartile range are shown. W3110, *n*=943; FW1541, *n*= 941. In B-E, measurements were performed in FW1541.

**B,** Cell area (left) and aspect ratio (right), respectively, plotted against length and width. n, population size; $R^2$, goodness of fit; r, Spearman coefficient; P: two-tailed probability.

**C,** Positive correlation between cell length and time based on experimental data acquired at 15 min intervals. Only data that cover the entire cell cycle were used. The best-fitting equation was used to calculate the length as a function of time.

**D**, The period-time correlation diagram converted according to Figure 1E and S3C.

**E,** Cell volume measurement of FW1541. n, population size; R2, goodness of fit; r, Spearman coefficient; P: two-tailed probability.

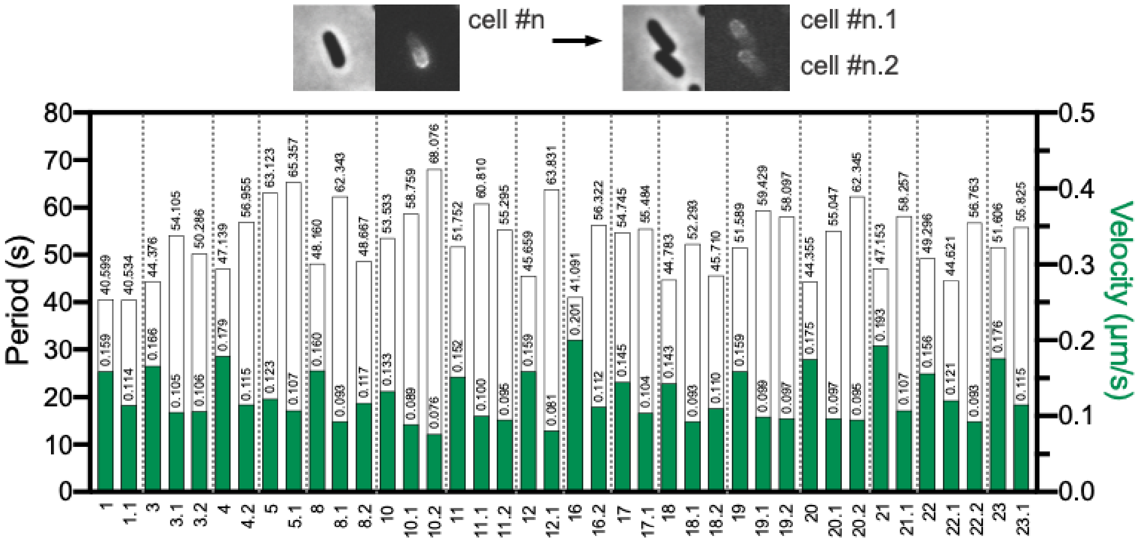


**Fig S4. Oscillation period and velocity before and after division suggest slowed oscillations in newborn cells.** Images were acquired at 15 min intervals over 2 hours. In some cases, the fluorescence of one of the two newborn cells was too weak to be studied. Numbers on the x-axis indicate cell lineages.

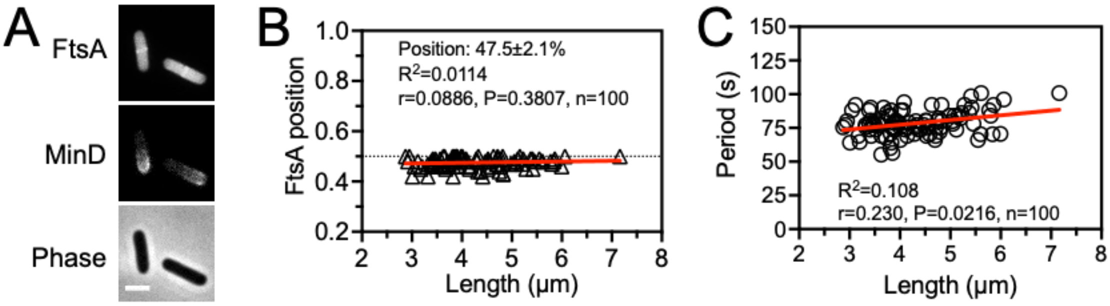


**Fig S5. Oscillation period, cell length and accuracy of division site placement.**

The ring-like structure of FtsA-mScarlet-I was located at a relative position of 47.5±2.1% (*n*=100) from either pole regardless of cell length, and period and velocity of the MinD oscillation. The cell center is set to position 0.5 for normalization of FtsA position. n, population size; R2, goodness of fit; r, Spearman coefficient; P: two-tailed probability.

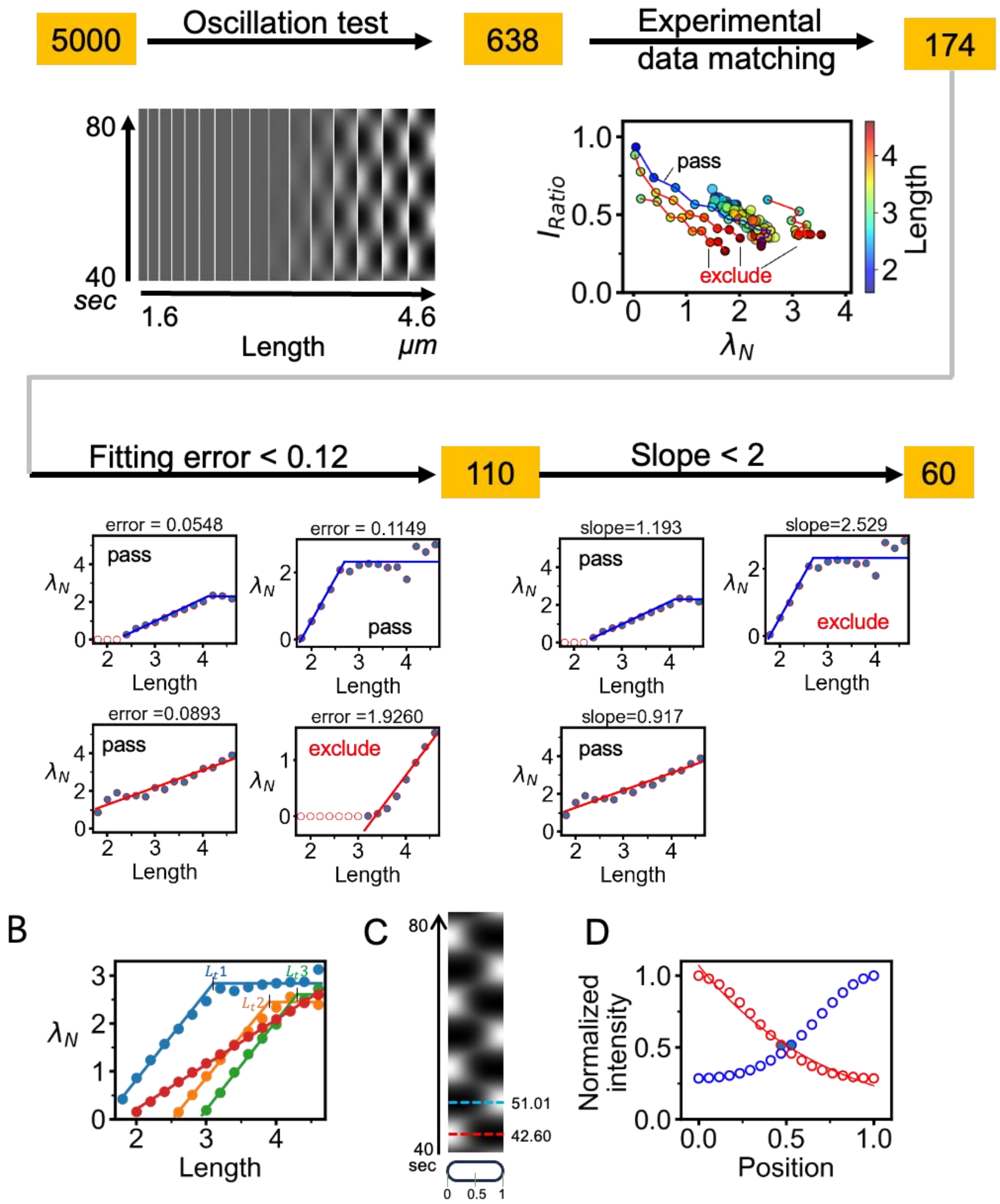


**Fig S6. Screening of rate constants and analysis of the simulation results.** This is a supplemental figure of Fig. 4.

**A**, Scheme of data processing and screening for combinations of rate constants that generate spatiotemporal oscillations.

**B**, Demonstration of the length-dependent change in $\lambda_N$ and the variable fitting error. Incremental $\lambda_N$ values are fitted with a linear equation $\lambda_N = aL + b$ or $\lambda_N$ = c, where $L$ is the length. While red lines show uniphasic fits, other colored lines show biphasic fits with different transition lengths $L_t$.

**C**, An example kymograph of a given length demonstrating oscillation of membrane-bound MinD ($C_d + C_{de}$) between 40 to 80 seconds. Concentrations are represented by intensities in simulations. The red and blue dotted lines mark the time points (42.60 and 51.01 s) when the maximum intensity at either pole is identified.

**D**, Determination of $I_{Ratio}$ at the intersection of the concentration profiles of the membrane-bound MinD at two cell halves. The left intensity profile (red hollow markers) is collected at 42.60 sec when the maximal intensity occurs. Each data point is normalized against the maximum intensity on this profile. Similarly, the right profile (blue hollow markers) is collected at 51.01 sec and processed. The resulting left and right profiles are symmetrical, with less than 0.02% difference between them. The two filled circles at the middle indicate the minimal relative concentration in this virtue cell, equivalent to $I_{Ratio}$. Here, the profile is fitted from the maximum to the minimum intensity using an exponential equation $y = 0.99\exp(-0.25x) + 0.064$, with $\lambda_N$ = 2.05.

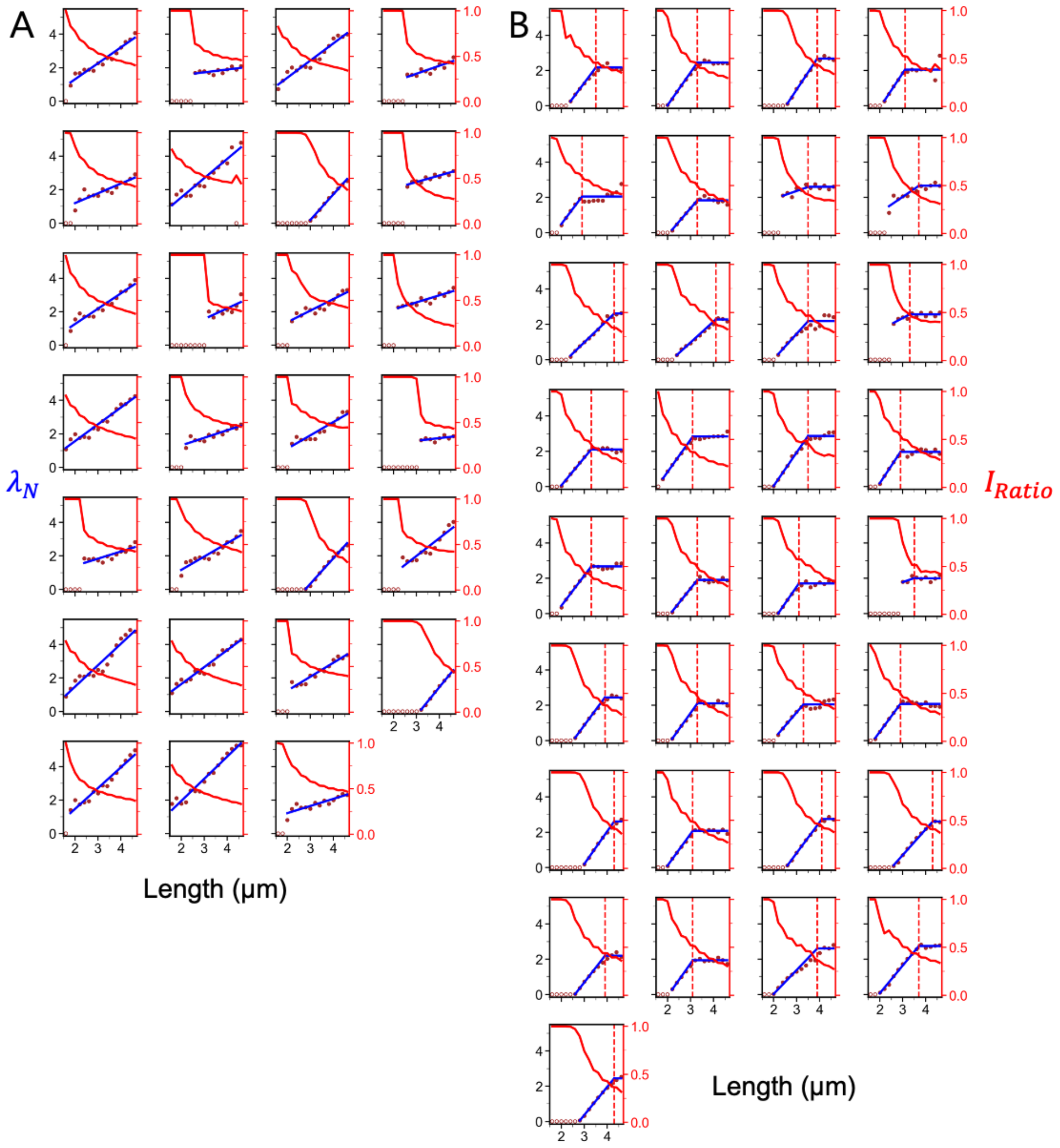


**Fig S7. Concentration profile analysis of simulated MinD dynamics,** demonstrated using normalized decay exponent $\lambda_N$ versus length (blue) and normalized $I_{Ratio}$ versus length (red). A, uniphasic group; B, biphasic group. The red dashed line in (B) marks the transition length $L_t$ from the incremental to plateau phase in the biphasic group. This figure is supplemental to Fig. 4 C, D.

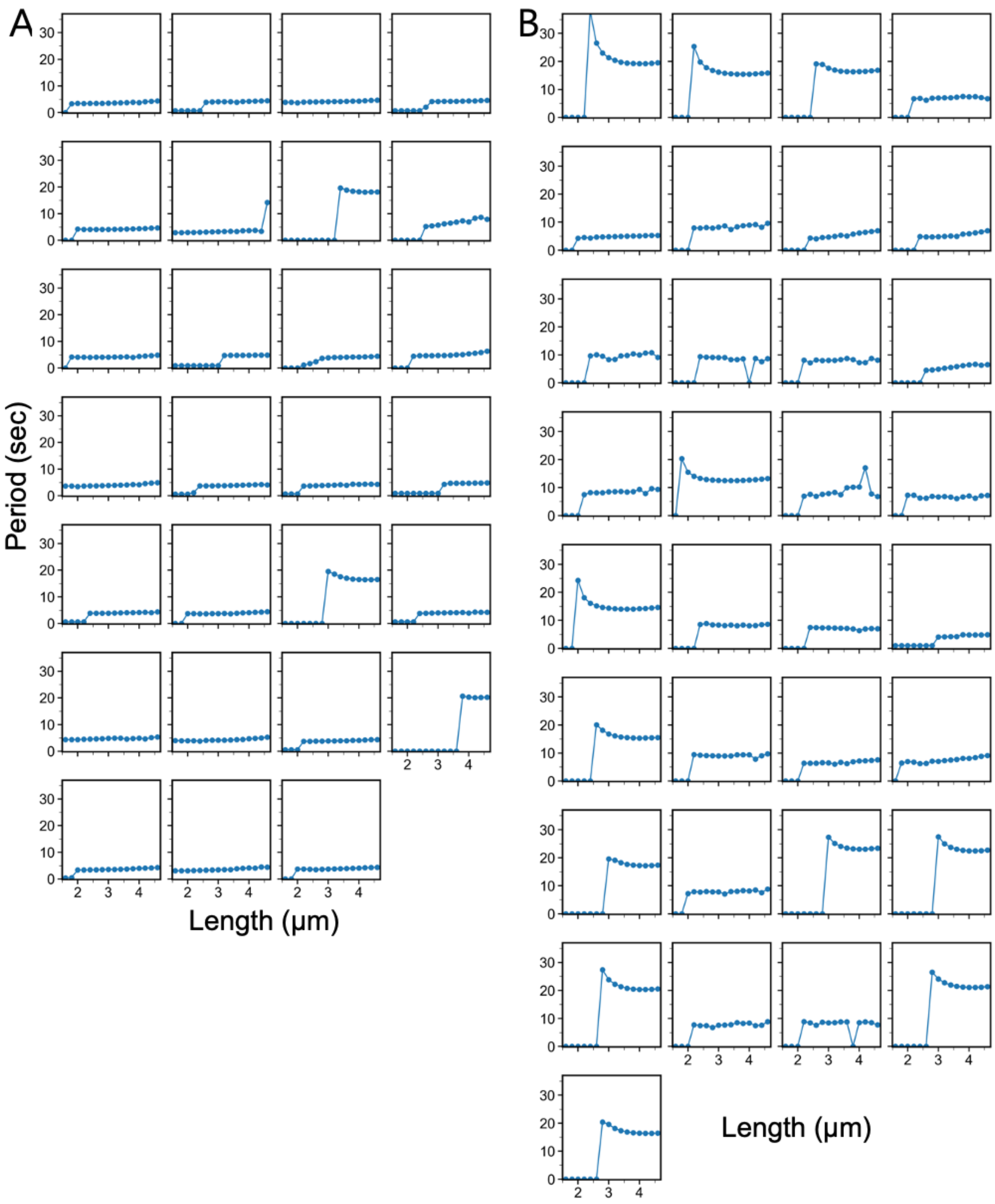


**Fig S8. Period analysis of simulated MinD dynamics**. A, uniphasic group; B, biphasic group. This figure is supplemental to Fig. 4B.

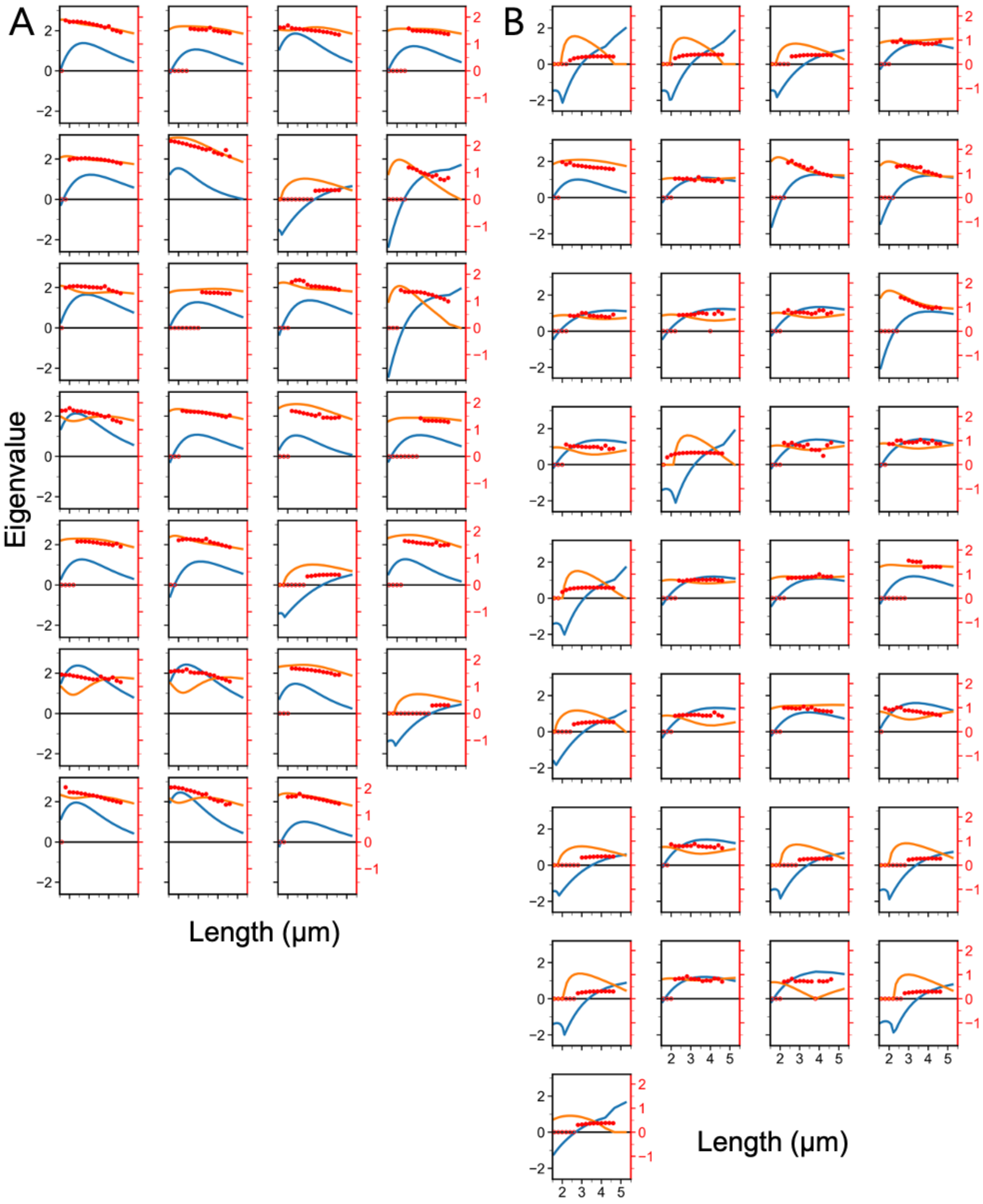


**Fig S9. Hopf instability plots showing the most divergent eigenvalue versus length**. These plots were created in a linear stability analysis to search for combinations of rate constants that cause spatiotemporal oscillations. Blue curve, real part; orange curve, imaginary part; red point, oscillation frequency. This is supplemental to Fig. 4B-D and S7,8.
